# Bayesian Inference of Gene Regulatory Networks at Stochastic Steady State

**DOI:** 10.64898/2026.01.10.698684

**Authors:** Anshi Gupta, Ryeongkyung Yoon, Krešimir Josić

**Affiliations:** Department of Mathematics, University of Houston, Houston, TX 77204, USA; Department of Biology and Biochemistry, University of Houston, Houston, TX 77204, USA; Department of Mathematics and Computer Science, Wabash College, Crawfordsville, IN 47933, USA; NSF-Simons National Institute for Theory and Mathematics in Biology, Chicago, IL 60611

## Abstract

Gene Regulatory Networks (GRNs) form the regulatory backbone that coordinates gene expression. The architecture of GRNs shapes their function and constraints the biochemical pathways through which information flows. Inferring the structure of regulatory interactions is thus essential for understanding biological systems, and designing targeted therapies. Despite substantial progress in GRN inference, most approaches – from statistical methods to deep learning – do not take into account fundamental biochemical processes that drive regulatory dynamics. To address this shortcoming, here we present a novel Bayesian inference approach based on using the Chemical Langevin Equation (CLE) as a model of gene expression dynamics at stochastic equilibrium. Interactions in GRNs are sparse, and we thus use a regularized horseshoe prior enabling selective shrinkage of unsupported interactions while identifying strong regulatory edges. We evaluate our method using synthetic gene expression data, allowing for benchmarking against a known ground truth. Our approach allows us to infer kinetic parameters, identify network structure, and infer regulatory cycles without the need to observe transient dynamics. This Bayesian alternative to current methods thus provides both biological interpretability and structural identifiability in GRN inference.

## 1 Introduction

Gene Regulatory Networks (GRNs) play a central role in controlling gene expression and coordinating cellular functions during development, cell differentiation, and responses to environmental stimuli [12]. Inferring the structure of GRNs is thus essential to understanding the regulatory mechanisms that govern complex biological processes. Despite extensive efforts to develop experimental and statistical approaches for GRN inference, reconstructing network structure and dynamics from gene expression data remains a challenging inverse problem. Key obstacles include stochasticity in gene expression [67], sparse connectivity between regulatory components [16], and nonlinear interactions within the network [45]. Sparse sampling and measurements near dynamical equilibrium further compound these challenges, making it difficult to resolve dynamic regulatory interactions from gene expression data.

Previous approaches have typically attempted to infer regulatory interactions from high-throughput, high-dimensional biological data, often without relying on detailed mechanistic assumptions. Techniques such as mutual information [46, 78], regression-based inference [36, 37], and deep learning [66] have been employed to recover GRN structure, aiming to balance predictive performance with interpretability. Although such methods capture complex, nonlinear gene interactions, issues with interpretability and validation persist. On the other hand, many model-based inference methods require measurements of transient dynamics: Since multiple parameter combinations are consistent with identical steady-state behavior, observations of transients are often needed to ensure parameter identifiability [35].

Few existing methods for GRN inference capture stochastic gene expression dynamics, enforce biologically motivated sparsity to infer network structure, interaction types, and kinetic parameters from data in stochastic equilibrium, while providing principled uncertainty quantification [38, 39]. To address this problem we introduce a Bayesian inference framework that describes intrinsic stochasticity using the Chemical Langevin Equation (CLE) [22, 30] and incorporates a regularized horseshoe prior [57] to reflect the sparsity of interactions in GRNs. We apply our method to synthetic data from GRNs at stochastic equilibrium, with measurements sampled from the stationary distribution. At equilibrium, the temporal structure of fluctuations provides information about regulatory interactions and kinetic parameters that is not captured by the moments of the distribution alone [61]. Our approach is closely related to the SINDy algorithm [8, 32, 43], and is conceptually related to fluctuation-dissipation relations in statistical physics, where equilibrium fluctuations reveal response properties without requiring external perturbations [77]. We show that we can identify the underlying network structure and infer kinetic parameters and interaction types, producing interpretable outputs with well-calibrated uncertainty estimates. Notably, we can recover features such as regulatory cycles and autoregulatory interactions, which are often missed by other Bayesian methods [20].

Our approach is based on a mechanistic model of regulatory interactions, in contrast to non-parametric models which offer greater flexibility but may sacrifice interpretability and biological grounding. We demonstrate reliable performance on small-scale networks, even when only partially observed. We thus lay the foundation for a robust, mechanistic network inference framework applicable in more complex settings.

## 2 Materials and Methods

Our goal is to infer the structure and regulatory parameters of gene regulatory networks (GRNs) from stationary gene expression data. To do so, we fit a mechanistic model of gene expression to data using a Bayesian approach. The resulting posterior distributions determine network topology (presence/absence of edges), interaction types (activation or repression), and reaction rates, while providing principled uncertainty quantification for all inferred quantities.

### 2.1 GRN model

A gene regulatory network (GRN) is a system in which regulatory molecules (primarily transcription factors (TFs), but also non-coding RNAs and other factors) regulate gene expression through activation or repression. Here, we concentrate on regulation by TFs, and represent a GRN as a directed graph with nodes corresponding to genes (or more precisely, gene products). For each gene, *i*, we denote by *X*_*i*_ (*t*) the abundance of the corresponding transcription factor at time *t*, and let X(*t*) = (*X*_1_(*t*), …, *X*_*p*_ (*t*)) be the state of the network, where *p* is the total number of TFs and genes. We assume reactions in a system at constant volume so that *X*_*i*_ (*t*) can represent the molecular count or concentration. A directed edge, *i* → *j*, represents a regulatory interaction where gene *i* (via its product) impacts the expression of gene *j*. Edges carry a sign indicating *activation* (the TF increases the target’s transcription rate) or *repression* (the TF decreases it). We allow for auto-regulation which is represented by a self-loop. At the molecular level, regulation arises when TFs bind promoter or enhancer regions and modulate RNA polymerase recruitment [56, 58]. Although other regulatory mechanisms are possible, here we consider only transcriptional regulation via activation and repression.

The relationship between TF concentrations and transcriptional output is nonlinear, due to saturation at high concentrations and regulatory thresholds [6, 41]. The dependence of transcription rate on TF concentration is commonly described using sigmoidal (Hill-type) functions, which approximate equilibrium TF–promoter binding and reproduce the nonlinear dose–response curves observed experimentally [3, 40, 62]. We model the effect of a single regulator on transcription rate by

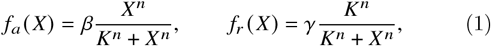

where *β* and *γ* denote the maximal transcription rates under activation (*f*_*a*_) and repression (*f*_*r*_), respectively. The parameter *K* is the half-maximal effective concentration, representing the TF level at which transcription reaches half of its maximal rate. The Hill coefficient, *n*, determines the steepness of the input–output curve and reflects the apparent cooperativity of the regulatory interactions. For simplicity, we assume negligible basal transcription, but our framework can be extended to include it.

Many genes are regulated by multiple TFs, and promoter activity often reflects their combined influence. Because the molecular mechanisms of combinatorial TF regulation are often unknown or intractable, promoter responses are typically modeled using coarse-grained logic functions including AND, OR, and hybrid gates [3, 9, 63, 73]. Here, we focus on the *AND gate*, and assume that transcription occurs only when all required TFs are bound to their respective binding sites. This represents a common regulatory motif with multiple inputs converging to activate transcription. While we only consider the case of AND gates with two TFs, the framework can be extended to other logic gates (OR, NOT, hybrid) and to genes with more than two regulators.

We use the following functions to describe AND gate type regulation for different combinations of two activators and repressors [55]:

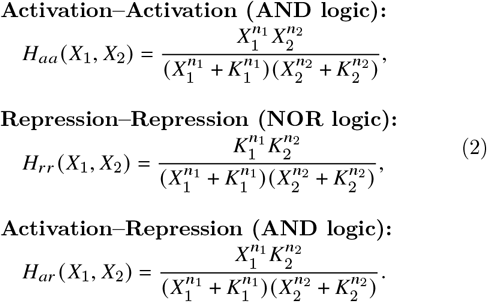

These normalized functions are multiplied by the maximal transcription rate to obtain the actual production rate.

Gene expression is inherently stochastic at the single-cell level due to the discrete nature of molecular events and the probabilistic timing of biochemical reactions. *Intrinsic noise* is due to the probabilistic nature of biochemical events (e.g., transcription factor or RNA polymerase binding), while *extrinsic noise* stems from cell-to-cell variability in global factors such as growth state and environmental influences [15, 54, 67, 68]. We focus on intrinsic noise as it directly reflects the biochemical processes we model (transcription, degradation, regulation), as well as the regulatory topology. Incorporating extrinsic noise would require including additional sources of variability.

We validate our approach using synthetic data. To do so we generate sample expression trajectories using the *stochastic simulation algorithm* (SSA), with the reaction rates as defined in Eqs. (1) and (2). The SSA produces trajectories whose probability law is specified by the *Chemical Master Equation* (CME) which determines the evolution of the probability distribution over all possible molecular states [22].

Direct inference using the CME quickly becomes computationally intractable. Instead, we use a discretized form (Euler– Maruyama) of the *Chemical Langevin Equation* (CLE) [23, 30]. The CLE is a mesoscopic SDE whose sample paths are distributed approximately according to the probability law described by the CME. Intuitively, the CLE replaces the discrete change in TF number over small time intervals by Gaussian increments with reaction propensities determining the mean and variance of each increment. This diffusion approximation is appropriate under well-stirred conditions, and is valid when molecule numbers are sufficiently large (typically > 50) so that continuous state variables are appropriate, yet small enough that stochastic fluctuations remain important. The discretized CLE yields tractable transition probabilities, enabling efficient likelihood-based parameter inference [25–27].

### 2.2 Example: Single gene autoregulation

As a first example, and to explain notation, we consider single-gene autoregulation [4, 42]. This simplest of GRNs can exhibit rich dynamics, and serves as the basis for multi-gene systems analyzed later. Let *X* denote the TF concentration, synthesized with a state-dependent propensity *f* (*X*) and degraded at rate *αX*, resulting in the following reactions

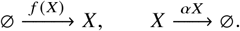

The corresponding CLE takes the form [30, 74]:

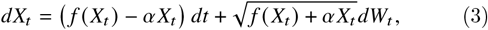

where *W*_*t*_ is a standard Wiener process. The drift term represents net synthesis and degradation, while the diffusion term accounts for stochastic fluctuations inherent in both processes. Setting *f* (*x*) equal to one of the Hill functions in Eqs. (1) allows us to model activation or repression.

The CLE framework provides a mechanistic description of stochastic gene expression, and can be used as a generative model for statistical inference of GRN structure and parameters given a sequence of TF measurements. When discretized using the Euler–Maruyama scheme, the conditional distribution of the state at time *X*_*t*+Δ*t*_ given the current state *X*_*t*_ is Gaussian, with its mean determined by the drift term and variance by the diffusion term [1, 8, 29, 64]. For parameter inference when *network architecture is known*, we can use the discretized version of Eq. (3) which takes the form

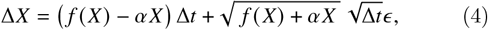

where ϵ ~ *N* (0, 1) denotes a sample from a standard Gaussian distribution, Δ*X* = *X* (*t*) − *X* (*t* − Δ*t*), and we suppressed the explicit dependence on *t*. This discretization yields a stochastic process, *X* (*t*), evolving in discrete time steps of size Δ*t* approximating the continuous in time process *X*_*t*_.

Our goal is also to infer the existence and type of interactions in a GRN. Thus we will not assume that the architecture of the GRN is known, but will use models that include a range of potential interactions. In the present example this means that for *inference* we use the following model which allows for both activation and repression,

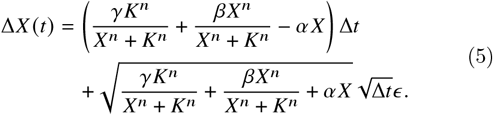

We assume that elements in the GRN either activate or repress one another, or do not interact. We discuss below how a Bayesian approach using Eq. (5) and horseshoe priors for *β* and *γ* results in successful inference with estimates 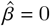 or 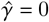, or both.

### 2.3 General Inference Model

Larger GRNs can be modeled by adding further nodes and interactions. The model we use for inference will always be a generalization of the model used to generate synthetic data that includes a range of potential interactions, *e*.*g*. compare Eq. (4) and Eq. (5). In the following we therefore only describe the model used for inference. We will see that a generalization of Eq. (5) yields explicit transition densities and can be used to define a likelihood function that can be evaluated directly from a sequence of TF measurements. The CLE that approximates the generative model described exactly by the CME also underlies the statistical model used for inference. Hence, the inferred parameters such as activation or repression rates, degradation constants retain their meaning in the underlying biochemical reactions.

To extend the inference model to larger GRNs, we represent activation, repression, and degradation using matrix-valued functions of the state vector. For simplicity, we suppress the explicit dependence on time in all models defined in this section. If the state of a GRN comprising *p* genes and the corresponding TFs is given by the vector ***X*** ∈ ℝ^*p*^, the discretized CLE generalizing Eq. (5) *in the absence of combinatorial regulation* has the form

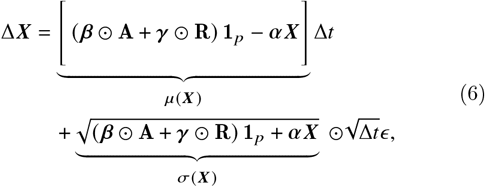

where **A** : ℝ^*p*^ → ℝ^*p*×*p*^ and **R** : ℝ^*p*^ → ℝ^*p*×*p*^ are matrix-valued functions. Each matrix element is a Hill function of a single state variable such that ***A***_*i, j*_ (***X***) = *f*_*a*_ (*X* _*j*_) captures the transcription rate of the TF encoded by gene *i* when regulated by the product of gene *j*. Similarly, *R*_*i j*_ (***X***) = *f*_*r*_ (*X* _*j*_) models repression. The matrix ***α*** ∈ ℝ^*p*×*p*^ is diagonal with protein-specific degradation/dilution rates, while ***β, γ*** ∈ ℝ^*p*×*p*^ contain the maximal transcription rates under activation and repression, respectively. Here and below the vector *ϵ* ∈ ℝ^*p*^ consists of *p* independent samples from the standard Gaussian distribution, and **1**_*p*_ ∈ ℝ^*p*^ is a vector with all entries equal to 1. In this and subsequent models, *μ*(***X***) and σ(***X***) denote the drift and diffusion terms, respectively.

For example, in a two gene regulatory network, the parameter matrices take the form

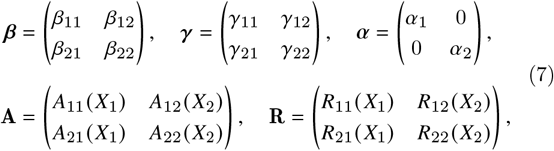

where the elements of the matrices **A** and **R** are Hill functions,

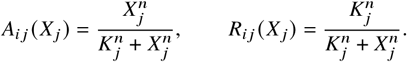

To capture combinatorial regulation, we extend the model to allow for two regulators per gene via an AND gate interaction:

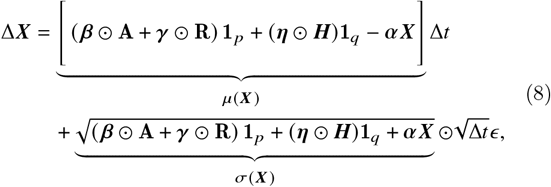

Here, *q* is the maximum number of regulatory input combinations that a gene can receive, and the matrix ***H*** : ℝ^*p*^ → ℝ^*p*×*q*^ contains Hill functions representing combinatorial regulation. Each entry 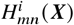 depends on two components of the state vector. For example, in a two-gene regulatory network with possible AND-gate regulation, *q* = 4 corresponding to the four possible two-input combinations so that the matrix ***H*** takes the form:

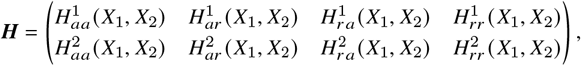

where the superscripts denote the gene index, and the subscripts, *a* and *r* indicate whether the corresponding regulator is activating or repressing. The functions used in this matrix are defined in Eq. (2). The parameter matrix ***η*** ∈ ℝ^*p*×*q*^ determines the strength of AND gate mediated regulation. The activation and repression matrices, ***A*** and ***R***, and the parameter matrices ***α, β, γ*** are defined as in the absence of combinatorial regulation, *i*.*e*. as in Eq. (6).

We found that Eq. (8) cannot be used to accurately infer parameters in GRNs with three or more nodes. Convergence and identifiability issues become more pronounced with an increasing number of horseshoe parameters (see Supplementary Fig. 10). To improve inference in the presence of combinatorial regulation, we thus performed inference in two steps. In the first step, we identify only the presence and absence of regulatory edges. We do so by introducing a matrix of Boolean variables, **Ω** ∈ {0, 1}^*p*×*q*^, whose elements indicate the presence or absence of AND gate mediated regulation, and matrix ***θ*** ∈ {0, 1}^*p*×*p*^ indicating whether regulatory interaction corresponds to single TF activation or repression. This reduces the number of parameters governed by the horseshoe prior, leading to faster and more stable inference. The resulting model is given by:

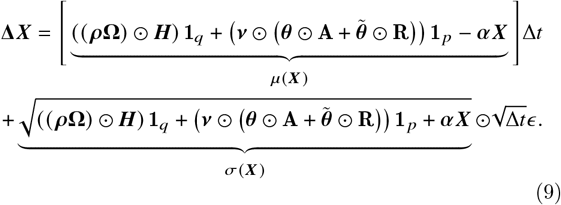

Here ***H, A*** and ***R*** are defined as in Eq. (8). Also, 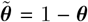 with the difference taken entry-wise. Thus *θ*_*i j*_ = 1 indicates the potential presence of an activation of gene *i* by the product of gene *j*, and the absence of corresponding repression. The matrix *ρ* ∈ ℝ^*p*×*p*^ determines the strength of the AND gate mediated regulation, while ***ν*** ∈ ℝ^*p*×*p*^ represents the strength of direct (single-input) regulatory interactions. The entries in row *i* of the matrix **Ω** determine the potential presence of AND gates at gene *i*: Let *ω*_*i, j*_ = 1 if the product of gene *j* activates gene *i* in combination with another TF, *ω*_*i, j*_ = 0 if it represses it, and let 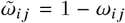 Then the elements in the *i*^th^ row of the matrix **Ω** are the *q* = 2*p*(*p* − 1) possible pairwise products of all combinations of *ω*_*i, j*_ and 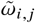 with differing indices *j*. Thus 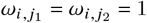 indicates potential activation-activation regulation of gene *i* by genes *j*_1_ and *j*_2_, while 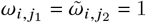 indicates potential activation-repression type regulation.

In the same example of two genes with possible AND-gate regulation discussed above, the matrix ***H*** remains unchanged, while the matrices *ρ*, **Ω, *ν***, and ***θ*** take the following forms:

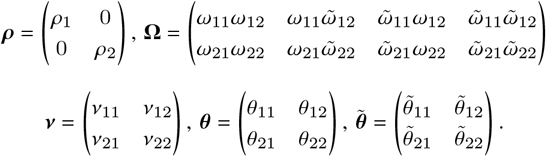

Here only one of the entries on each row of **Ω** is non-zero, since only one of the possible interactions listed in Eq. (2) can be present.

In the second inference step we infer the reaction rates. To do so we use the GRN architecture inferred in the first step, and the model given by Eq. (8). Assuming a fixed GRN architecture allows us to infer the reaction rates by estimating a substantially reduced number of parameters.

Thus, we use Eq. (6) for inference under the assumption that each gene has a single regulator. Eq. (9) is a generalization that we use to allow for possible combinatorial regulation. In the former case, inference requires estimating only continuous parameters (***β, γ, α***), while in the latter both continuous (*ρ*, ***ν, α***) and discrete (***θ*, Ω**) parameters need to be estimated. These descriptions of the evolution of gene products allow us to define appropriate likelihoods, and infer both network structure and kinetic rates from data. We describe this inference framework next.

### 2.4 Bayesian inference framework

We next show how to use Bayesian inference to infer both GRN network architecture and reaction rates from data [7]. Bayes’ Theorem relates model parameters, *θ*, and the observed measurements, 𝒟, via the relation *p* (*θ* | 𝒟) ∝ *p* (𝒟 | *θ*) *p* (*θ*). In the present case the likelihood *p* (𝒟 | *θ*) is determined by the discretized CLE. Using the Euler–Maruyama approximation, the increments, Δ*X*, follow a Gaussian distribution with mean determined by the drift term and variance by the diffusion term in the CLE [8]. The likelihood of a full trajectory is then the product of Gaussians evaluated at the observed counts or concentrations across all time steps,

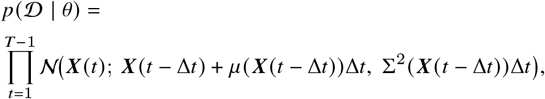

where 𝒩 (***x***; ***μ*, Σ**) denotes the probability density function of a Gaussian distribution with mean ***μ*** and covariance **Σ** evaluated at ***x***. Here, the variance matrix is diagonal and equals Σ^2^(***X***) = diag(σ^2^(***X***)), where σ^2^(***X***) is a vector obtained by squaring each component of the diffusion vector σ(***X***) elementwise. The variance matrix is diagonal because in the underlying biochemical reaction model the noise terms for different species arise from independent reaction channels [23]. Thus the increments follow the laws defined by the generative models in Eqs. (6)–(9).

#### Parameters Estimated

In all examples, we estimated only the maximal transcription rates and degradation rates, which were consistently identifiable from the data. In contrast, the parameters ***K*** (half-maximal concentration) and *n* (Hill coefficient) were generally more difficult to infer. For instance, as shown in Fig. 1(e), when input concentrations fall within the saturated region of the response curve, the system becomes insensitive to changes in ***K***, rendering this parameter difficult to identify. In contrast, Fig. 1(f) illustrates that when the input spans the dynamic range around ***K***, its influence on the output becomes observable, enabling more reliable inference. Even in such regimes, inference of ***K*** remained sensitive to prior assumptions; good estimates required using informative priors. The Hill coefficient, *n*, was typically even more weakly identifiable. It is known that ***K*** and *n* are often structurally or practically unidentifiable, as they do not strongly determine the system’s dynamics [11, 31, 48]. We thus fixed both parameters to their known values during inference.

**Figure 1.**
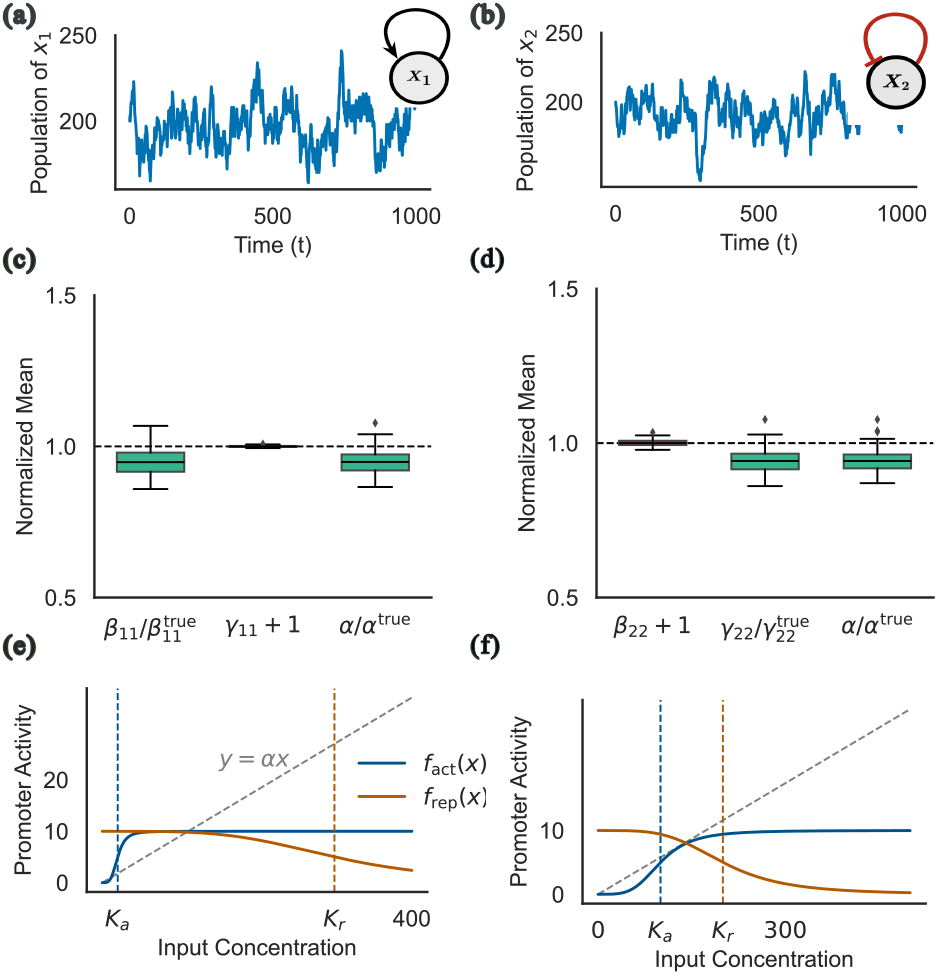
Inference of regulation in a single-gene system. **(a)-(b)** TF time-series and the corresponding network structures for a single gene: (a) self-activating, and (b) self-repressing regulatory network. Each realization was generated using the Gillespie algorithm. Parameters were chosen so that the marginal means and variances of the simulated trajectories were approximately equal for both systems. **(c)-(d)** Normalized boxplots of the posterior means of the inferred reaction parameters for the self-activating (c) and self-repressing (d) models across 50 realizations. Here, and in the following figures, we show the ratio between the posterior mean and the true value when the true value is nonzero (green), whereas the posterior mean of parameters whose true value is zero is shifted by 1 (red). Hence, estimates close to 1 indicate accurate inference. **(e)-(f)** Activation (blue) and repression (red) Hill function in the identifiable (f) and non-identifiable (e) parameter regimes. Detailed results and summary statistics are provided in the Supplementary Material.

#### Priors

We used uninformative priors in all examples. These priors could be changed to include available information about the structure of the GRN or reaction rates. We distinguish two cases corresponding to the two model formulations:

- *Single-regulator model (no combinatorial regulation) (Eq. (6))*. Reaction rates are determined by the activation strengths, ***β***, repression strengths, ***γ***, and degradation rates, ***α***. To promote sparsity in interactions, we used horseshoe priors for ***β*** and ***γ*** [57]. These priors reflect the need for strong evidence for an interaction, and the assumption that a TF cannot both activate and repress the same target, *i*.*e*. that for each gene pair, (*i, j*), at most one of *β*_*i j*_ = 0 and *γ*_*i j*_ = 0 is nonzero. These shrinkage priors strongly pull most interaction strengths toward zero while allowing a subset to remain large, consistent with the expectation that only a few of all possible interactions are present in GRNs.
- *Combinatorial model (Eq. (9))*. In addition to single-input activation and repression regulatory strengths in the parameter matrix ***ν*** and degradation rates ***α***, this model includes the gate-strength matrix, *ρ*, as well as matrix of binary indicator variables, ***θ***, corresponding to activation/repression and **Ω** corresponding to AND gate regulation. We used regularized horseshoe priors for parameters ***ρ*** and ***ν*** to reflect the assumption of sparsity in both gated and non-gated interactions. The binary indicators are assigned independent, uninformative Bernoulli(0.5) priors, representing the prior belief that each potential interaction or gate is equally likely to exist or not.

For the degradation rates, ***α***, which are always positive, we used uninformative half-normal priors. This enforces non-negativity while constraining the inferred values to the expected range of degradation rates.

To improve sampling efficiency and avoid issues related to thick Cauchy tails when using horseshoe priors, we adopted the recommendations by Piironen and Vehtari [57] and replaced the Half-Cauchy priors with half-Student-*t* distributions.

Specifically, we set

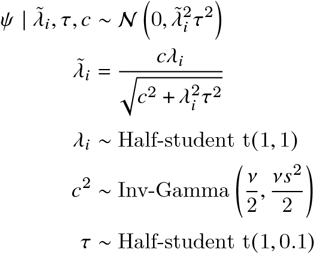

Here *ψ* represents parameters for which we used the horse-shoe prior. Following standard practice we chose *v* = 4 and *s* = 2 for the inverse-gamma prior on *c*^2^. This formulation provides robust shrinkage, ensures well-behaved posteriors, and facilitates efficient posterior sampling for high-dimensional or weakly identified models [32, 57].

#### Approximating the posterior distributions

Samples from the posterior distributions were generated using PyMC [2, 53]. For models with only continuous parameters, we used Hamiltonian Monte Carlo (HMC) [5] with the No-U-Turn Sampler (NUTS) [33], accelerated via the BlackJAX backend for efficient sampling on CPUs and GPUs. When discrete parameters were present (e.g. indicator matrices), we reverted to PyMC’s standard NUTS implementation, which marginalizes over discrete variables when possible. Convergence was assessed using standard diagnostics, including the 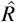 statistic, effective sample size, and visual inspection of trace plots.

### 2.5 Synthetic data

We generated synthetic gene-expression trajectories using the stochastic simulation algorithm (SSA) [22, 24]. In all simulations, we discarded an initial transient and retained measurements after the system equilibrated. In this regime molecular counts fluctuated randomly, but with stable statistical properties.

We chose the sampling interval Δ*t* to be sufficiently small so that the assumed discretization provides a good approximation of the underlying dynamics, and sufficiently large so that increments in TF numbers are approximately Gaussian and the correlation between consecutive observations was not excessive. For each parameter setting, we generated 50 independent realizations, each consisting of 3, 000 measurements. Replicate datasets were generated to ensure robust inference and allowed us to account for variability across stochastic sample paths [68]. All SSA simulations were implemented using the Python package GillesPy2 [47].

## 3 Results

GRNs govern cellular behavior through transcriptional regulation, with transcription factors (TFs) activating or repressing target genes. Inferring GRN structure and dynamics from gene expression data is challenging due to the complexity of regulatory interactions and the stochasticity of gene expression. We show that the proposed Bayesian inference framework allows us to simultaneously recover network topology, regulatory modes (activation/repression), and kinetic parameters from time-series expression data.

We demonstrate the performance of the proposed inference method using synthetic data from a range of canonical gene regulatory motifs. Throughout, we consider GRNs at stochastic equilibrium, with molecular fluctuations having reached their stationary probability distribution. ODE and logic-based models cannot capture stationary fluctuations in molecule numbers [71]. Structural inference of GRNs using such models therefore requires the observation of transients. In contrast, we show that a generative model that accounts for intrinsic stochasticity allows for the inference of kinetic parameters and regulatory structure from measurements of GRNs at stochastic equilibrium.

We illustrate the proposed method using a sequence of examples of increasing complexity. We begin with single-gene autoregulation, then consider the classic two-gene toggle switch [21], followed by an extension to a toggle with combinatorial regulation. We next consider the three-gene repressilator [14], a canonical oscillatory circuit, and finally turn to a coherent type-1 feed-forward loop [3, 44]. This allows us to assess both the accuracy and the scalability of the framework.

Throughout we work in rescaled (non-dimensional) time, so all parameters are expressed without physical units. Under this convention, the transcription rate parameters correspond to the maximal expected number of molecules transcribed per unit rescaled time. In all examples we performed inference on 50 independent realizations to quantify robustness. We also report posterior means (denoted by 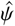 for a parameter *ψ*) and highest density intervals (HDIs) for a single, representative realization. We show representative results here, while the full inference results along with a complete description of the models used for inference are provided in the Supplementary Material.

### 3.1 Single-Gene Motifs

We first consider a single-gene autoregulatory circuit. In this motif, the product of a gene regulates its own production, as shown in Fig. 1(a,b). Here, and in all subsequent examples, black edges represent activation and red edges repression. We consider both self-activation and self-repression, with transcription rates defined by Hill functions given in Eq. (1). Details about the parameters used to generate the synthetic data for this and subsequent examples are provided in the Supplementary Material.

We first asked whether we can use the proposed inference framework to distinguish between activation and repression from trajectories in stochastic equilibrium. To do so, we generated synthetic TF trajectories for both cases using parameters that produced marginal distributions with matched means and variances. Example trajectories are shown in Fig. 1(a,b). The associated marginal distributions of TF counts are nearly indistinguishable (see Supplementary Fig. 1). Many existing inference approaches, such as moment-matching [61] and fitting to the steady-state distribution [28], use only marginal statistics and would therefore fail to distinguish activation and repression from such data.

Despite the similarity in marginal distributions, we successfully inferred the correct regulatory mode in all 50 independent realizations. In the activation case, the repression parameter *γ*_11_ was estimated to be near zero (posterior mean 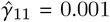, HDI: [−0.95, 1.05]), whereas the activation parameter *β*_11_ was inferred to be close to its true value of 10 (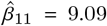, HDI: [8.31, 9.87]). Conversely, in the repression case, the activation parameter *β*_22_ was inferred to be near zero (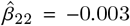, HDI: [−1.02, 0.94]), while the repression parameter *γ*_22_ was estimated to be close to 10 (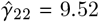, HDI: [8.74, 10.40]) (see Supplementary Table 3 for exact values). Fig. 1(c) and (d) show the normalized box plots of the inferred parameters for self-activation and self-repression across 50 independent SSA realizations.

Key to the successful discrimination of regulatory modes are the observation of the temporal dynamics: Self-activation and self-repression exhibit distinct autocorrelation structures and transient responses to fluctuations, even when their steady-state distributions are identical [13]. Using the discretized CLE as a generative model exploits these temporal features to accurately infer regulatory modes that are difficult to distinguish using marginal TF distributions alone.

However, we found that the system is not identifiable in the *saturated regimes* of the Hill function (Fig. 1(e)). At sufficiently high or very low concentrations of the transcription factor, the Hill function saturates and fluctuations in counts of a regulating TF have negligible effect on the production rate. In this regime, the likelihood function is approximately constant, making it difficult to distinguish between activation and repression, and infer the transcription rates. This limitation is well recognized: in saturated regimes, different Hill functions can produce indistinguishable observable statistics [62]. A likelihood-based inference framework helps to determine when such identifiability issues arise [60], and to determine the parameter regimes in which the likelihood is sufficiently sensitive to changes in TF count to make inference possible (Fig. 1(f)). In all the examples discussed in this text, we therefore work with synthetic data generated in this regime.

### 3.2 Two-Gene Motifs

We next consider a two gene toggle-switch–type network, in which one gene activates a partner while being repressed in return as shown in Fig. 2(a). Such canonical motifs have been studied in both natural regulatory circuits and synthetic gene networks [19, 76]. We generated synthetic data from the GRN shown in Fig. 2(a) and modeled the system using Eq. (6), where the two-gene case is provided as an illustrative example (see Eq. (7) for the corresponding matrices). For inference, we used a fully connected GRN model that encompasses all potential interactions (See Supplementary Eq (2) for the definition of the likelihoods).

**Figure 2.**
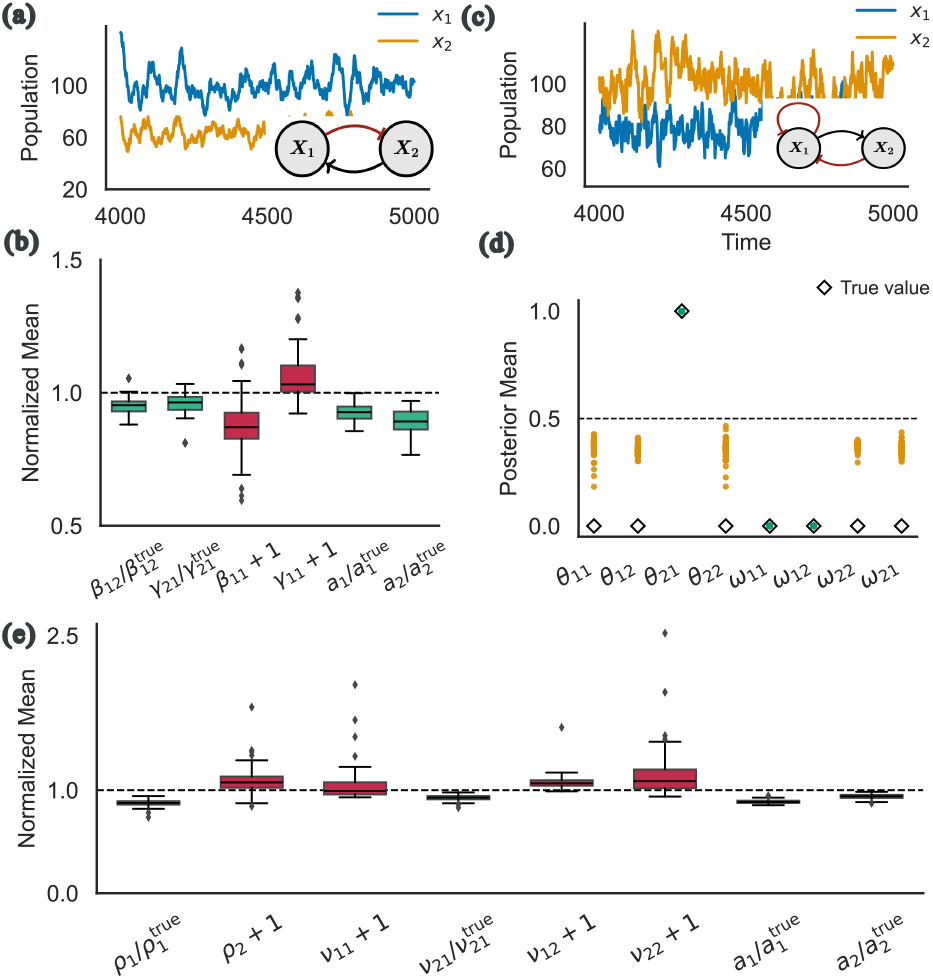
Inference in two-gene regulatory networks with and without combinatorial regulation. (a) Expression levels and corresponding network structures for a two-gene system without combinatorial regulation. (b) Normalized boxplots of the posterior means of the inferred parameters obtained from 50 realizations of the trajectories. (c) Expression levels for a two-gene regulatory system with combinatorial AND-gate regulation. (d) Posterior means of the inferred binary regulatory indicators (*θ, ω*) obtained from 50 time series realizations. We used a threshold of 0.5 to determine the presence and type of regulatory interactions. (e) Normalized posterior means of the inferred reaction rates. Here and in subsequent figures we present results for a representative subset of key parameters; detailed results are provided in the Supplementary Material.

We again accurately recovered the underlying regulatory network in 50 independent realizations. The activation parameter *β*_12_ (true value = 10) was inferred to be close to its true value 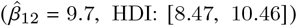, correctly identifying gene 2 as an activator of gene 1. Likewise, the repression parameter *γ*_21_ (true value = 20) was estimated to be close to 20(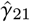, HDI: [17.05, 21.31]), accurately capturing the repressive interaction. The degradation parameters *α*_1_ and *α*_2_ were also accurately estimated (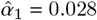, HDI: [0.022, 0.032] and 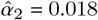, HDI: [0.011, 0.022], respectively), corresponding to relative errors below 5%. All other regulatory parameters *β*_11_ and *γ*_11_ corresponding to nonexistent interactions had posterior means below 0.2, with 95% HDIs restricted to values near zero. Hence, we did not identify any spurious interactions.

These inference results are summarized in Fig. 2(b), which demonstrates that we can accurately infer both the structure of the GRN and the parameters governing TF dynamics at a stochastic equilibrium.

### 3.3 Two-Gene motifs with an AND gate

We next considered the case of a two-gene motif with both cross-regulation and autoregulation shown in Fig. 2(c). Such network motifs, combining cross-feedback with autoregulation, are common in both natural and synthetic gene networks [34] and can generate rich dynamical behaviors including oscillations and bistability [10, 52]. We again focus on the non-oscillatory regime.

This circuit poses a stringent test for network inference because autoregulatory interactions can be obscured by cross-regulatory signals and because self-regulation produces temporal correlations that can confound parameter estimation [72]. Successful recovery of both auto-and cross-regulatory interactions in this motif thus demonstrates the ability to disentangle multiple regulatory modes acting on the same gene.

We used synthetic trajectories from a system in which gene 1 was regulated by gene 2 and by its own product shown in Fig. 2(c). Regulation was modeled using AND-gate logic according to Eq. (9). For inference, we used Eq. (9) instead of Eq. (8) because it is computationally more efficient, requiring the inference of fewer parameters.

For inference, we again used a fully connected GRN model that included all potential interactions, including autoregulation (See supplementary Eq (4)). Binary-valued variables in the matrices **Ω** and **Θ** indicate whether potential regulatory interactions were activating or repressive, while continuous-valued parameters (*ρ, ν*) quantified the interaction strength (See Section 2.3). When a continuous parameter was inferred to be close to zero (*i*.*e*. when the corresponding HDI is concentrated around 0), the corresponding edge was assumed to be absent. Thus, edge existence and interaction strengths were inferred simultaneously within a single inference step.

We were able to infer the correct two-gene AND gate motif structure from all 50 independent synthetic trajectories. The scatter plots in Fig. 2(d) and Fig. 2(e) show the posterior means for the binary variables and continuous variables, respectively. In this example, the presence of regulatory interactions is determined by the product of the corresponding binary and continuous variables, specifically through the terms *θ*_*i j*_ *ν*_*i j*_ and *ρ*_*i*_*ω*_*i j*_ *ω*_*ik*_, which becomes negligible when either component is inferred to be close to zero. In Fig. 2(d), orange dots represent binary variables associated with parameters whose true value are zero.

Since gene 1 represses its own production and is repressed by *X*_2_, its reaction is described by a repression-repression term, 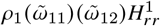. Green dots in Fig. 2(d) correspond to 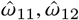 being close to zero indicating the presence of combinatorial AND gate repression at gene 1. The parameter *ρ*_1_ is inferred to be close to its true value of 20 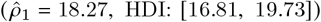, while the parameters *θ*_11_ and *θ*_12_ are estimated to be close to zero (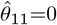 and 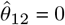).The corresponding parameters, *ν*_11_ and *ν*_12,_ are also inferred to be close to zero (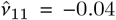, HDI: [−0.61, 0.48]) and (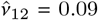, HDI: [−0.76, 1.02]) (See Fig. 2(e)), indicating the absence of the corresponding edges in the GRN.

Gene 2 transcription is activated by the product of gene 1, which is captured by the term *ν*_21_*θ*_21_ ***A***_21_. As shown in Fig. 2(d) and (e), green dots correspond to the estimate 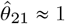. The related parameter *ν*_21_ indicates a strong activating effect of gene 1 on gene 2 (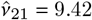, HDI: [8.82, 10.03]). On the other hand, the parameter *ν*_22_ (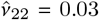, HDI: [−0.65, 0.78]) and *ρ*_2_ (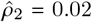, HDI: [−0.84, 0.94]) was inferred to be close to zero, indicating the absence of self-regulation and combinatorial regulation at gene 2.

Thus, the inferred GRN structure and parameters were consistent with the motif structure we used to generate the data, and we were able to infer both the existence of combinatorial regulation and the strengths of the underlying interactions in the presence of multiple regulatory inputs.

### 3.4 Repressilator

We next considered the repressilator [50], a synthetic circuit that allows us to test the performance of the proposed inference methods with cyclic regulatory architectures. The repressilator consists of three transcriptional repressors arranged in a cyclic inhibitory loop) [14], as shown in Fig. 3(a). While this network can generate sustained oscillations, we focused on the non-oscillatory regime, where trajectories converge to a stochastic equilibrium.

**Figure 3.**
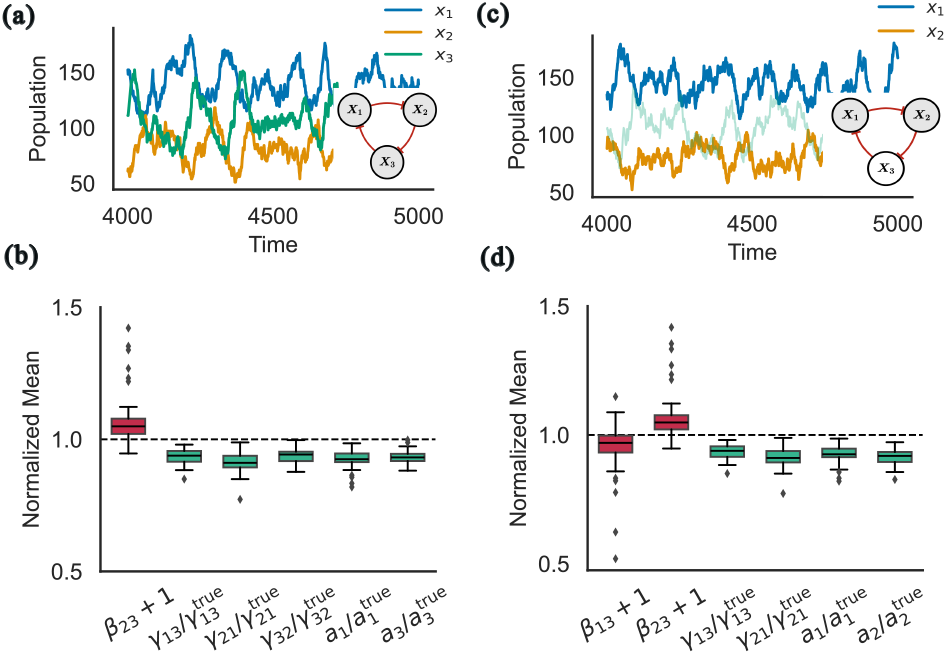
Inference in the non-oscillatory repressilator. (a) Trajectories of TF levels in a fully observed repressilator system (*X*_1_ ⊣ *X*_2_ ⊣ *X*_3_ ⊣ *X*_1_). (b) Normalized boxplots of the posterior means of inferred regulatory parameters obtained from 50 realizations of the trajectories. (c) The same repressilator system when only partially observed with the product of gene 3 (light green line) not used for inference. (d) The normalized boxplots of the inferred parameters show that we recovered the correct regulatory topology even when the GRN is only partially observed.

We were again able to both identify the correct regulatory structure, and accurately estimate the maximal transcription rates, as well as degradation rates from synthetic data. The boxplots in Fig. 3(b) show that the recovered parameters closely match the ground truth. Across 50 independent realizations, the repression parameters *γ*_13_, *γ*_21_ and *γ*_32_ were inferred with posterior mean estimates close to their true values of 10 (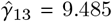, HDI:[8.43, 10.36]), 20 (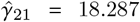, HDI:[16.53, 19.86]), and 15 (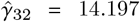, HDI:[13.20, 15.15]), respectively. All other interaction parameters (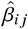 and 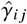) were estimated to be close to zero, correctly indicating the absence of other interactions. The degradation rates,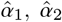 and 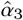,were also recovered accurately.

### 3.5 Partial Observability

In practice, we can often observe only a subset of a GRN. This poses significant challenges for network inference and parameter identifiability [69]. To evaluate our method’s performance under partial observability, we returned to the three-gene repressilator motif but assumed that only the products of genes 1 and 2 are measurable (Fig. 3(c)). This allowed us to test what interactions can be inferred when the network is partially observed.

Despite the absence of direct measurements of the product of gene 3, we recovered all parameters governing the dynamics of the *observed* nodes. We were able to infer that gene 1 represses gene 2, with *γ*_21_ estimated close to its true value of 20 (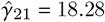, HDI:[16.46, 19.84]), and that gene 3 represses gene 1, with *γ*_13_ estimated near its true value of 10 (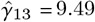, HDI:[8.43, 10.34]). The degradation rates were also inferred accurately, with 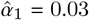, HDI:[0.027, 0.032] and 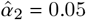, HDI:[0.042, 0.050]. All other parameters governing the evolution of *X*_1_ and *X*_2_ were also estimated well. The horseshoe prior effectively suppressed spurious interactions, with the estimated activation parameters *β*_13_ and *β*_23_ taking posterior means near zero (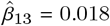, HDI:[−0.29, 0.38]) and (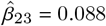, HDI:[−0.24, 0.62]), respectively, indicating the absence of the corresponding regulatory effects.

As expected, parameters determining the dynamics of products of the unobserved gene 3, including 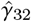 and 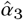, had large posterior uncertainty. However, the parameter *γ*_13_ captures the regulatory impact of the unobserved gene 3 on the observed system dynamics, and was inferred correctly. Thus, our method correctly inferred the presence and strength of this hidden regulatory link without generating false positive interactions, demonstrating robust performance under partial observability.

### 3.5 Coherent Feedforward Loop (C1-FFL)

The coherent type-1 feedforward loop (C1-FFL) is one of the most frequently occurring regulatory motifs in transcriptional networks [3, 44]. It consists of a regulator that activates a downstream target both directly and indirectly through an intermediate regulator, with all three interactions being activating. C1-FFLs generate *sign-sensitive delays*, in which activation of the target requires sustained input. This property enables the motif to filter out transient fluctuations and contributes to the robustness of gene expression [44].

To model this GRN we assumed that gene 1 is transcribed at a constant rate, and activates the expression of gene 2, while both the products of gene 1 and 2 jointly activate the expression of gene 3 as shown in Fig. 4(a). In the model used for inference we did not allow for self loops as they significantly. increased the number of possible combinations of interactions at each gene. Although they could be incorporated, doing so would considerably increase computational demands. We performed inference in two steps since inferring both the GRN structure and transcription rates produced unreliable results. These difficulties were due in part to limitations of the Hamiltonian Monte Carlo (HMC/NUTS) samplers, which frequently exhibited poor chain mixing and persistent divergences, often caused by chains becoming trapped. Sampling performance improved substantially even over the course of this project as the HMC/NUTS implementations in PyMC were refined (see Conclusion).

**Figure 4.**
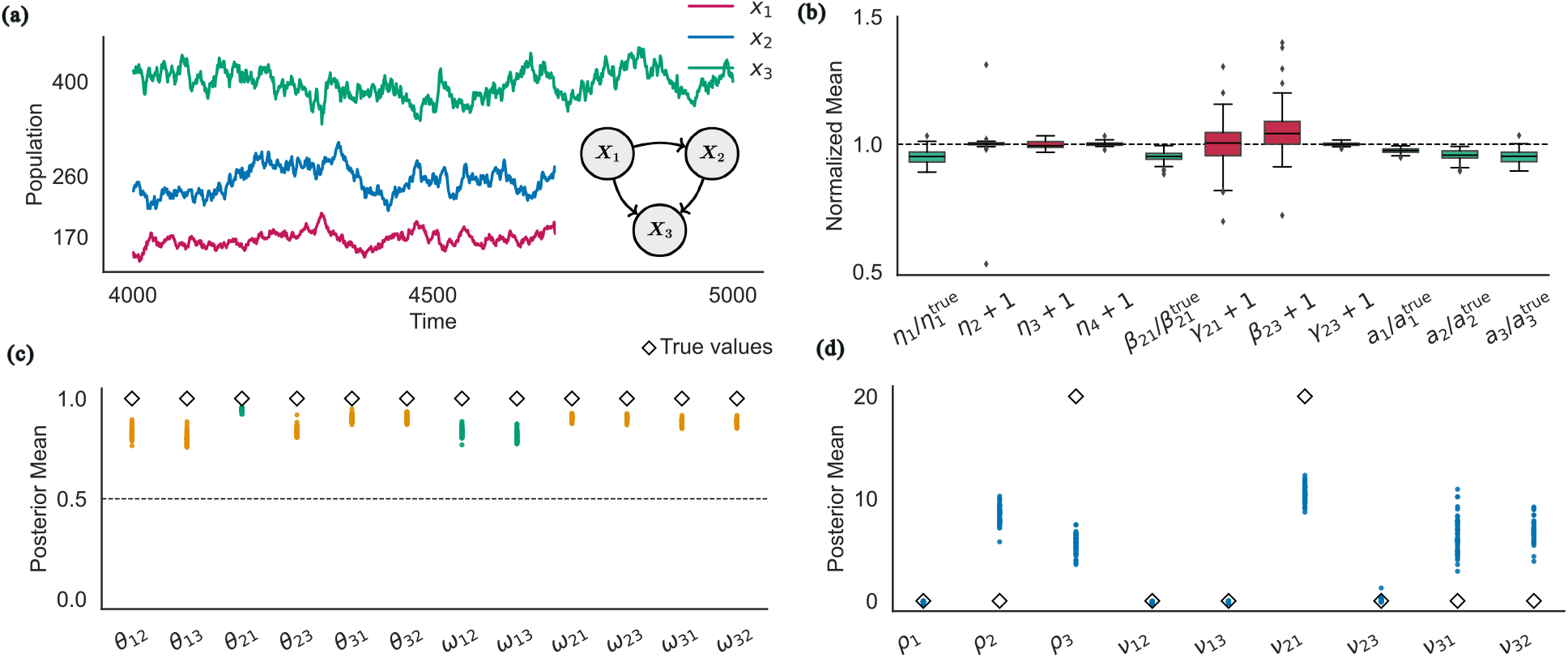
Results of the two-step inference procedure applied to the coherent type-1 feed-forward loop (C1-FFL) motif. (a) Gene Expression trajectories generated of a network motif where gene *X*_1_ is produced at a constant rate of 5 units, activates *X*_2_, and *X*_3_ is jointly activated by both *X*_1_ and *X*_2_ through an AND gate. (b) Normalized boxplots from the second inference step, showing the ratios of posterior means to true parameter values for *β* (activation strength), *α* (degradation rate), and *η* (combinatorial regulation strength). (c) Scatter plot of inferred binary regulatory indicators (*θ, ω*) from the first inference step, where a threshold of 0.5 was used to determine the presence and type of regulatory interactions. (d) Scatter plots of the inferred continuous parameters (*ν, ρ*) from the first inference step, representing the strength of individual and combinatorial regulatory interactions.

The first inference step was focused on inferring GRN architecture. Here we used Eq. (9) to define the likelihood of the model parameters. This formulation of the generative model incorporates binary indicator variables to identify active regulatory interactions. When the associated continuous parameters had HDIs concentrated around zero, the corresponding interactions were assumed to be absent. Pruning these potential interactions yielded a reduced model that we used in the second inference step focused on inferring interaction strengths and degradation rates. We used Eq. (8), with the architecture inferred in the first step. This two-stage approach reduced the effective dimensionality of the parameter space and mitigated the influence of spurious edges on rate estimates.

In the first step across 50 realizations the model consistently inferred 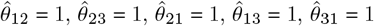, and 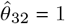, indicating that all six potential interactions were present (See Fig. 4(c)). The AND-gate binary indicator variables *ω*_*i j*_ were also inferred to be active: 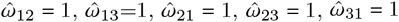, and 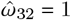.

As shown in Fig. 4(d), the AND-gate indicators were estimated 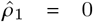, HDI:[−0.71, 0.36] and 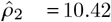, HDI:[−0.36, 19.33], suggesting no AND-gate regulation at gene 1 but a potential combinatorial effect at gene 2. The single-input parameters acting on gene 2 were inferred as 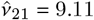, HDI:[−0.27, 19.33] and 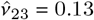, HDI:[−0.52, 0.78]. Since only 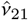 admits clearly nonzero values, we concluded that *X*_1_ regulates *X*_2_, ruling out the presence of an actual AND gate at gene 2.

For gene 3, *ρ*_3_ was inferred to be nonzero (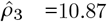, HDI:[−0.40, 19.75]), and the corresponding single-input parameters, estimated as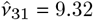, HDI:[−0.46, 19.42] and 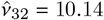, HDI:[−0.34, 19.76], had wide HDIs that included substantial nonzero values, indicating an AND-gate interaction involving TFs *X*_1_ and *X*_2_. In the second step, only the 11 AND gate regulation of *X*_3_ was retained, with the single-input parameters *ν*_31_ and *ν*_32_ pruned to zero. Although the single-input terms had positive estimated values, they were pruned because the strong interaction term *ρ*_3_ alone was sufficient to explain the data. This resulted in a simpler and more interpretable model, in which gene 3 is activated exclusively by the simultaneous presence of both transcription factors *X*_1_ and *X*_2_, with no independent activation by either factor alone.

The final inference results demonstrate that the structure of the GRN and the interaction strengths were accurately recovered (See Fig. 4(b)). Thus, a two step approach consisting of first identifying the GRN structure, and then estimating the corresponding rate parameters can improve inference in complex networks, enabling both accurate identification of network architecture and parameter estimation.

## 4 Discussion

We have shown that the Chemical Langevin Equation (CLE) provides a tractable and biologically grounded framework for inferring network structure and kinetic parameters in gene regulatory networks operating in the mesoscopic regime, where intrinsic stochasticity is significant but molecule counts are sufficiently large to justify a continuous approximation. The CLE captures both deterministic reaction kinetics and stochastic fluctuations within a unified stochastic differential equation framework, making it well-suited for modeling gene expression, where noise plays important functional roles [15, 59]. While exact likelihood-based inference using the Chemical Master Equation (CME) is theoretically optimal, it becomes computationally intractable for multi-gene networks [51]. The CLE offers a practical alternative that retains essential stochastic information while remaining amenable to efficient Bayesian inference algorithms. The CLE assumes a well-stirred, constant-volume system and does not provide a good approximation when molecule numbers approach zero [23]. Despite these limitations, our results demonstrate that a CLE-based inference approach can be used to successfully recover network topology and parameters from simulated gene expression time series across a range of biologically relevant regimes, including systems with cross-regulation and strong regulatory feedback.

Central to the proposed Bayesian approach is the use of horseshoe priors [32] which reflect the assumed sparsity of interactions, and allow for the accurate recovery of network architecture. We have shown that such priors suppress spurious edges while retaining true regulatory interactions. We have also tried alternatives, such as Laplacian priors [65] (not shown), and found that well tuned horseshoe priors provided consistently sharper discrimination between existing and absent interactions.

We used a particular form of regulatory interactions to define the statistical model and the resulting likelihood functions, such as the AND gate model of cross-regulation. Other regulatory logic functions can be incorporated in this framework, and our approach can be extended to determine which function best fits the data if multiple candidates are included in the model.

Inference in gene regulatory networks is subject to both structural uncertainty – the inability to distinguish between architectures – and parameter identifiability when either the equations do not uniquely determine certain parameters regardless of how much data is available [60], or, in practice, when finite, noisy observations do not constrain parameter estimates [17, 70]. We consistently recovered the correct network topology in all test cases. However, we observed practical identifiability limitations for certain parameters. Specifically, in the case of a coherent type-1 feed-forward network, during the first step of the inference procedure, structural identifiability issues became evident. Some parameters were linearly correlated; for example, *ν*_21_ and *ρ*_2_ were consistently assigned similar values, indicating that the model could not uniquely distinguish them. Similarly, *ρ*_2_ and *ν*_23_, *ν*_32_ and *ρ*_3_, and *ν*_31_ and *ρ*_3_ also showed strong correlations. These dependencies limit the accuracy with which individual parameter values can be inferred. These correlation patterns are shown in Supplementary Fig. 10.

Our approach has several limitations. First, computational and data requirements grow with network size: inference for *N* genes involves *O*(*N*^2^) parameters, making inference in larger networks (> 5 genes) challenging. Second, for validation we used only synthetic data under idealized conditions. While this is a necessary first step in validating an inference method, we ignored extrinsic noise, measurement error, and parameter non-stationarity present in real experiments [15]. Third, our model assumes Hill function kinetics, CLE approximations valid at moderate copy numbers, and negligible time delays – assumptions that can be violated in biological systems and may thus impact inference. Future work should address these limitations through benchmarking, experimental validation, and model extensions that incorporate biologically important system properties.

Several avenues are available to address these challenges. First, incorporating both intrinsic and extrinsic noise sources into the likelihoods would yield more biologically realistic models. Second, systematic validation on experimental time-series data will be essential to establish practical utility. Third, improving scalability through approximate inference schemes [75], dimensionality reduction [49], or variational Bayesian approaches [18] will broaden applicability to larger systems. Finally, extending the framework to more complex network architectures, including feedback loops and cross-regulatory modules, could help uncover design principles of natural gene networks at larger scales. Together, these directions can enhance both the computational feasibility and biological relevance of the method, paving the way for systems-level studies of gene regulation.

## Acknowledgements

This work was supported by funding from the joint National Science Foundation and National Institutes of Health grant 1R01GM144959 and National Science Foundation EFRI-OI grant 2515431 (A.G. and K.J.).

## Code Availability

Code implementing the proposed inference algorithm and reproducing all results in the text is available at http://bit.ly/3Yq14Rx

## References

[1] A. Aalto, L. Viitasaari, P. Ilmonen, L. Mombaerts, and J. Gonçalves. Gene regulatory network inference from sparsely sampled noisy data. Nature Communications, 11(1):3493, 2020.

[2] O. Abril-Pla, V. Andreani, C. Carroll, L. Dong, C. J. Fonnesbeck, M. Kochurov, R. Kumar, J. Lao, C. C. Luhmann, O. A. Martin, et al. Pymc: a modern, and comprehensive probabilistic programming framework in Python. PeerJ Computer Science, 9:e1516, 2023.

[3] U. Alon. An Introduction to Systems Biology: Design Principles of Biological Circuits. Chapman and Hall/CRC, 2nd edition, 2019.

[4] A. Becskei, B. Séraphin, and L. Serrano. Positive feedback in eukaryotic gene networks: cell differentiation by graded to binary response conversion. The EMBO Journal, 20(10):2528–2535, 2001.

[5] M. Betancourt. A conceptual introduction to Hamiltonian Monte Carlo. arXiv preprint 1701.02434, 2017. Preprint, available at 1701.02434.

[6] L. Bintu, N. E. Buchler, H. G. Garcia, U. Gerland, T. Hwa, J. Kondev, and R. Phillips. Transcriptional regulation by the numbers: models. Current Opinion in Genetics & Development, 15(2):116–124, 2005.

[7] G. E. P. Box and G. C. Tiao. Bayesian Inference in Statistical Analysis. John Wiley & Sons, 1973. Reprinted 2011.

[8] S. L. Brunton, J. L. Proctor, and J. N. Kutz. Sparse identification of nonlinear dynamics with control (SINDYc). IFAC-PapersOnLine, 49(18):710–715, 2016.

[9] N. E. Buchler, U. Gerland, and T. Hwa. On schemes of combinatorial transcription logic. Proceedings of the National Academy of Sciences, 100(9):5136–5141, 2003.

[10] Y. Chen, J. K. Kim, A. J. Hirning, K. Josić, and M. R. Bennett. Emergent genetic oscillations in a synthetic microbial consortium. Science, 349(6251):986–989, 2015.

[11] O.-T. Chis, J. R. Banga, and E. Balsa-Canto. Structural identifiability of systems biology models: a critical comparison of methods. PLoS ONE, 6(11):e27755, 2011.

[12] E. H. Davidson and M. Levine. Gene regulatory networks. Proceedings of the National Academy of Sciences, 102(14):4935–4936, 2005.

[13] M. J. Dunlop, R. S. III Cox, J. H. Levine, R. M. Murray, and M. B. Elowitz. Regulatory activity revealed by dynamic correlations in gene expression noise. Nature Genetics, 40(12):1493–1498, 2008.

[14] M. B. Elowitz and S. Leibler. A synthetic oscillatory network of transcriptional regulators. Nature, 403(6767):335– 338, 2000.

[15] M. B. Elowitz, A. J. Levine, E. D. Siggia, and P. S. Swain. Stochastic gene expression in a single cell. Science, 297(5584):1183–1186, 2002.

[16] C. Espinosa-Soto. On the role of sparseness in the evolution of modularity in gene regulatory networks. PLoS Computational Biology, 14(5):e1006172, 2018.

[17] C. E. FitzGerald, S. Reich, V. Agaba, A. Mathur, M. S. Werner, and N. M. Mangan. Practical indistinguishability in a gene regulatory network inference problem, a case study. arXiv preprint 2508.21006, 2025. Preprint, available at 2508.21006.

[18] C. W. Fox and S. J. Roberts. A tutorial on variational Bayesian inference. Artificial Intelligence Review, 38(2):85–95, 2012.

[19] P. François and V. Hakim. Core genetic module: the mixed feedback loop. Physical Review E, 72(3):031908, 2005.

[20] N. Friedman, M. Linial, I. Nachman, and D. Pe’er. Using Bayesian networks to analyze expression data. Journal of Computational Biology, 7(3-4):601–620, 2000.

[21] T. S. Gardner, C. R. Cantor, and J. J. Collins. Construction of a genetic toggle switch in Escherichia coli. Nature, 403(6767):339–342, 2000.

[22] D. T. Gillespie. Exact stochastic simulation of coupled chemical reactions. The Journal of Physical Chemistry, 81(25):2340–2361, 1977.

[23] D. T. Gillespie. The chemical langevin equation. The Journal of Chemical Physics, 113(1):297–306, 2000.

[24] D. T. Gillespie. Stochastic simulation of chemical kinetics. Annual Review of Physical Chemistry, 58:35–55, 2007.

[25] A. Golightly and D. J. Wilkinson. Bayesian inference for stochastic kinetic models using a diffusion approximation. Biometrics, 61(3):781–788, 2005.

[26] A. Golightly and D. J. Wilkinson. Bayesian sequential inference for stochastic kinetic biochemical network models. Journal of Computational Biology, 13(3):838–851, 2006.

[27] A. Golightly and D. J. Wilkinson. Bayesian parameter inference for stochastic biochemical network models using particle Markov chain Monte Carlo. Interface Focus, 1(6):807–820, 2011.

[28] U. Herbach, A. Bonnaffoux, T. Espinasse, and O. Gandrillon. Inferring gene regulatory networks from single-cell data: a mechanistic approach. BMC Systems Biology, 11(1):105, 2017.

[29] D. J. Higham. An algorithmic introduction to numerical simulation of stochastic differential equations. SIAM Review, 43(3):525–546, 2001.

[30] D. J. Higham. Modeling and simulating chemical reactions. SIAM Review, 50(2):347–368, 2008.

[31] K. E. Hines, T. R. Middendorf, and R. W. Aldrich. Determination of parameter identifiability in nonlinear biophysical models: a Bayesian approach. Journal of General Physiology, 143(3):401–416, 2014.

[32] S. M. Hirsh, D. A. Barajas-Solano, and J. N. Kutz. Sparsifying priors for Bayesian uncertainty quantification in model discovery. Royal Society Open Science, 9(2):211823, 2022.

[33] M. D. Hoffman and A. Gelman. The No-U-Turn sampler: adaptively setting path lengths in Hamiltonian Monte Carlo. Journal of Machine Learning Research, 15(1):1593–1623, 2014. Available at http://jmlr.org/papers/v15/hoffman14a.html.

[34] S. Huang, Y.-P. Guo, G. May, and T. Enver. Bifurcation dynamics in lineage-commitment in bipotent progenitor cells. Developmental Biology, 305(2):695–713, 2007.

[35] X.-N. Huang, W.-J. Shi, Z. Zhou, and X.-J. Zhang. The identifiability of gene regulatory networks: the role of observation data. Journal of Biological Physics, 48(1):93– 110, 2022.

[36] V. A. Huynh-Thu and P. Geurts. dyngenie3: dynamical genie3 for the inference of gene networks from time series expression data. Scientific Reports, 8(1):3384, 2018.

[37] V. A. Huynh-Thu, A. Irrthum, L. Wehenkel, and P. Geurts. Inferring regulatory networks from expression data using tree-based methods. PLoS ONE, 5(9):e12776, 2010.

[38] G. M. James, C. Sabatti, N. Zhou, and J. Zhu. Sparse regulatory networks. The Annals of Applied Statistics, 4(2):663–686, 2010.

[39] R. Jiang, P. Singh, F. Wrede, A. Hellander, and L. Petzold. Identification of dynamic mass-action biochemical reaction networks using sparse Bayesian methods. PLoS Computational Biology, 18(1):e1009830, 2022.

[40] H. Kim and E. Gelenbe. Stochastic gene expression modeling with Hill function for switch-like gene responses. IEEE/ACM Transactions on Computational Biology and Bioinformatics, 9(4):973–979, 2012.

[41] T. Kuhlman, Z. Zhang, M. H. Jr. Saier, and T. Hwa. Combinatorial transcriptional control of the lactose operon of Escherichia coli. Proceedings of the National Academy of Sciences, 104(14):6043–6048, 2007.

[42] H. Kuwahara and O. S. Soyer. Bistability in feedback circuits as a byproduct of evolution of evolvability. Molecular Systems Biology, 8:564, 2012.

[43] N. M. Mangan, S. L. Brunton, J. L. Proctor, and J. N. Kutz. Inferring biological networks by sparse identification of nonlinear dynamics. IEEE Transactions on Molecular, Biological and Multi-Scale Communications, 2(1):52–63, 2016.

[44] S. Mangan, A. Zaslaver, and U. Alon. The coherent feedforward loop serves as a sign-sensitive delay element in transcription networks. Journal of Molecular Biology, 334(2):197–204, 2003.

[45] S. Manicka, K. Johnson, M. Levin, and D. Murrugarra. The nonlinearity of regulation in biological networks. NPJ Systems Biology and Applications, 9(1):10, 2023.

[46] A. A. Margolin, I. Nemenman, K. Basso, C. Wiggins, G. Stolovitzky, R. Dalla Favera, and A. Califano. Aracne: an algorithm for the reconstruction of gene regulatory networks in a mammalian cellular context. BMC Bioinformatics, 7(Suppl 1):S7, 2006.

[47] S. Matthew, F. Carter, J. Cooper, M. Dippel, E. Green, S. Hodges, M. Kidwell, D. Nickerson, B. Rumsey, J. Reeve, L. R. Petzold, K. R. Sanft, and B. Drawert. GillesPy2: a biochemical modeling framework for simulation driven biological discovery. Letters in Biomathematics, 10(1):87– 103, 2023.

[48] T. R. Middendorf and R. W. Aldrich. The structure of binding curves and practical identifiability of equilibrium ligand-binding parameters. Journal of General Physiology, 149(1):121–147, 2017.

[49] K. R. Moon, D. van Dijk, Z. Wang, S. Gigante, D. B. Burkhardt, W. S. Chen, K. Yim, A. van den Elzen, M. J. Hirn, R. R. Coifman, et al. Visualizing structure and transitions in high-dimensional biological data. Nature Biotechnology, 37(12):1482–1492, 2019.

[50] S. Müller, J. Hofbauer, L. Endler, C. Flamm, S. Widder, and P. Schuster. A generalized model of the repressilator. Journal of Mathematical Biology, 53(6):905–937, 2006.

[51] B. Munsky and M. Khammash. The finite state projection algorithm for the solution of the chemical master equation. The Journal of Chemical Physics, 124(4):044104, 2006.

[52] J. Panovska-Griffiths, K. M. Page, and J. Briscoe. A gene regulatory motif that generates oscillatory or multiway switch outputs. Journal of The Royal Society Interface, 10(79):20120826, 2013.

[53] A. Patil, D. Huard, and C. J. Fonnesbeck. Pymc: Bayesian stochastic modelling in Python. Journal of Statistical Software, 35(4):1–81, 2010.

[54] J. Paulsson. Summing up the noise in gene networks. Nature, 427(6973):415–418, 2004.

[55] R. Phillips, J. Bois, and M. Meyer. Analysis of feed-forward loops. http://be150.caltech.edu/2020/content/lessons/05_ffls.html, 2020. Caltech BE/Bi 150: Biological Circuit Design, Lesson 5.

[56] R. Phillips, J. Kondev, J. Theriot, and H. G. Garcia. Physical Biology of the Cell. Garland Science, 2 edition, 2012.

[57] J. Piironen and A. Vehtari. Sparsity information and regularization in the horseshoe and other shrinkage priors. Electronic Journal of Statistics, 11(2):5018–5051, 2017.

[58] M. Ptashne and A. Gann. Genes & Signals. Cold Spring Harbor Laboratory Press, 2002.

[59] A. Raj and A. van Oudenaarden. Nature, nurture, or chance: stochastic gene expression and its consequences. Cell, 135(2):216–226, 2008.

[60] A. Raue, C. Kreutz, T. Maiwald, J. Bachmann, M. Schilling, U. Klingmüller, and J. Timmer. Structural and practical identifiability analysis of partially observed dynamical models by exploiting the profile likelihood. Bioinformatics, 25(15):1923–1929, 2009.

[61] J. Ruess and J. Lygeros. Moment-based methods for parameter inference and experiment design for stochastic biochemical reaction networks. ACM Transactions on Modeling and Computer Simulation, 25(2):8, 2015.

[62] M. Santillán. On the use of the Hill functions in mathematical models of gene regulatory networks. Mathematical Modelling of Natural Phenomena, 3(2):85–97, 2008.

[63] T. Schlitt and A. Brazma. Current approaches to gene regulatory network modelling. BMC Bioinformatics, 8(Suppl 6):S9, 2007.

[64] D. Schnoerr, G. Sanguinetti, and R. Grima. Approximation and inference methods for stochastic biochemical kinetics—a tutorial review. Journal of Physics A: Mathematical and Theoretical, 50(9):093001, 2017.

[65] M. Seeger, F. Steinke, and K. Tsuda. Bayesian inference and optimal design in the sparse linear model. In Proceedings of the 11th International Conference on Artificial Intelligence and Statistics (AISTATS), pages 444–451, 2007. Available at http://proceedings.mlr.press/v2/seeger07a.html.

[66] H. Shu, J. Zhou, Q. Lian, H. Li, D. Zhao, J. Zeng, and J. Ma. Modeling gene regulatory networks using neural network architectures. Nature Computational Science, 1(7):491–501, 2021.

[67] P. S. Swain, M. B. Elowitz, and E. D. Siggia. Intrinsic and extrinsic contributions to stochasticity in gene expression. Proceedings of the National Academy of Sciences, 99(20):12795–12800, 2002.

[68] Y. Taniguchi, P. J. Choi, G.-W. Li, H. Chen, M. Babu, J. Hearn, A. Emili, and X. S. Xie. Quantifying E. coli proteome and transcriptome with single-molecule sensitivity in single cells. Science, 329(5991):533–538, 2010.

[69] A. F. Villaverde. Observability and structural identifiability of nonlinear biological systems. Complexity, 2019:8497093, 2019.

[70] A. F. Villaverde, A. Barreiro, and A. Papachristodoulou. Structural identifiability of dynamic systems biology models. PLoS Computational Biology, 12(10):e1005153, 2016.

[71] R.-S. Wang, A. Saadatpour, and R. Albert. Boolean modeling in systems biology: an overview of methodology and applications. Physical Biology, 9(5):055001, 2012.

[72] Y. Wang and S. He. Inference on autoregulation in gene expression with variance-to-mean ratio. Journal of Mathematical Biology, 86(5):87, 2023.

[73] A. Warmflash and A. R. Dinner. Signatures of combinatorial regulation in intrinsic biological noise. Proceedings of the National Academy of Sciences, 105(45):17262–17267, 2008.

[74] D. J. Wilkinson. Stochastic modelling for quantitative description of heterogeneous biological systems. Nature Reviews Genetics, 10(2):122–133, 2009.

[75] D. J. Wilkinson. Stochastic Modelling for Systems Biology. Chapman and Hall/CRC, 3rd edition, 2018.

[76] K. Williams, M. A. Savageau, and R. M. Blumenthal. A bistable hysteretic switch in an activator–repressor regulated restriction–modification system. Nucleic Acids Research, 41(12):6045–6057, 2013.

[77] C.-C. S. Yan and C.-P. Hsu. The fluctuation-dissipation theorem for stochastic kinetics—implications on genetic regulations. The Journal of Chemical Physics, 139(22):224109, 2013.

[78] X. Zhang, X.-M. Zhao, K. He, L. Lu, Y. Cao, J. Liu, J.-K. Hao, Z.-P. Liu, and L. Chen. Inferring gene regulatory networks from gene expression data by path consistency algorithm based on conditional mutual information. Bioinformatics, 28(1):98–104, 2012.

