## Supplementary Material for "Bayesian Inference of Gene Regulatory Networks at Stochastic Steady State"

This section provides detailed descriptions of the synthetic datasets, likelihood formulations, and expanded inference results for each gene–regulatory motif described in the main text. For every motif, we include:

- **Data generation:** Synthetic trajectories generated using the stochastic simulation algorithm (SSA), together with the propensity functions and the true parameter values used in each simulation. These are summarized in tables specific to each motif.
- **Likelihood specification:** The drift and diffusion terms entering the Gaussian transition density obtained from the chemical Langevin equation (CLE).
- **Expanded inference results:** Posterior summaries (mean, standard deviation, and $\hat{R}$ statistics) for all inferred parameters, together with joint posterior distributions and box plots.

Each section below follows this structure consistently: (i) description of the synthetic data, (ii) explicit drift and diffusion terms for the likelihood, and (iii) expanded posterior results including all diagnostic figures and tables.

### 1 Notation Table

We summarize all notations and model parameters used in this paper in the notation table below.

Table 1: Key notation used in the Section 2

| Symbol | Meaning |
|---|---|
| $p$ | Number of genes/TFs in the network |
| $\mathbf{X}(t) \in \mathbb{R}^p$ | State vector of TF concentrations |
| $\boldsymbol{\beta}, \boldsymbol{\gamma} \in \mathbb{R}^{p\times p}$ | Activation and repression strength matrices |
| $\boldsymbol{\alpha} \in \mathbb{R}^{p\times p}$ | Diagonal matrix of degradation rates |
| $\mathbf{A}(\mathbf{X}), \mathbf{R}(\mathbf{X}) \in \mathbb{R}^{p\times p}$ | Matrix-valued Hill functions for activation/repression |
| $q$ | Number of possible combinatorial input combinations |
| $\boldsymbol{\rho} \in \mathbb{R}^{p\times q}$ | AND-gate regulatory strengths |
| $\boldsymbol{\Omega} \in \{0,1\}^{p\times q}$ | Binary indicators for AND gates |
| $\boldsymbol{\nu} \in \mathbb{R}^{p\times p}$ | Single-reaction regulatory strengths |
| $\boldsymbol{\theta} \in \{0,1\}^{p\times p}$ | Binary indicators for activation vs. repression |

### 2 Single-Gene Motifs

#### 2.1 Data

We generate synthetic time-series data using the stochastic simulation algorithm (SSA). The regulatory functions and parameter values used to simulate single-gene systems are summarised in Table 2. Parameters were chosen so that their marginal statistics remain nearly the same (See histograms shown in Fig. 1). We chose $\Delta t = 1$ to generate the synthetic trajectories.

As shown in Fig. 2, this sampling interval strikes a balance between temporal resolution and autocorrelation decay, and in our case was found to be the optimal choice. The same value of $\Delta t$ was used to generate all synthetic datasets across the examples presented in this work.

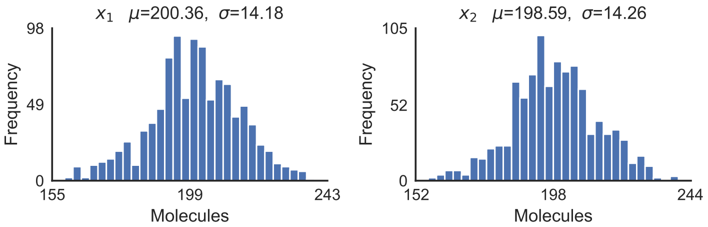


Figure 1: Histograms of the time series for the single gene self-activating (left) and self-repressing system (right). Both exhibit nearly identical marginal means and variances.

Table 2: Propensity functions and parameter values used to generate the SSA data for the single-gene motifs.

| Transcription Factor | Propensities | $\beta/\gamma$ | $K$ | $n$ | $\alpha$ |
|---|---|---|---|---|---|
| $X_1$ | $f_a(X_1),\ \alpha_1 X_1$ | 10 | 29 | 4 | 0.05 |
| $X_2$ | $f_r(X_2),\ \alpha_2 X_2$ | 10 | 500 | 4 | 0.05 |

#### 2.2 Definition of likelihoods

The drift and diffusion terms used to define the likelihood in the single gene case are:

$$\begin{aligned} \mu_i &= \left( \frac{\beta_{ii}\, X_i^n}{X_i^n + K_i^n} + \frac{\gamma_{ii}\, K_i^n}{X_i^n + K_i^n} - \alpha_i\, X_i \right) \Delta t, \\ \sigma_i &= \left( \sqrt{\frac{\beta_{ii}\, X_i^n}{X_i^n + K_i^n} + \frac{\gamma_{ii}\, K_i^n}{X_i^n + K_i^n} + \alpha_i\, X_i} \right) \sqrt{\Delta t}. \end{aligned} \tag{1}$$

The likelihood for each species is then modeled using a normal transition density $N(\mu_i, \sigma_i)$ for $i = 1, 2$.

#### 2.3 Expanded Results for Inference

Table 3 reports the posterior summaries for all inferred parameters. Across all chains, the $\hat{R}$ values were equal to 1, indicating excellent convergence and well-mixed samples. The posterior means closely matched the true values used to generate the data, demonstrating accurate recovery of the model parameters. The joint posterior distributions, along with the marginal posterior distributions on the diagonal, are shown in Fig. 3.

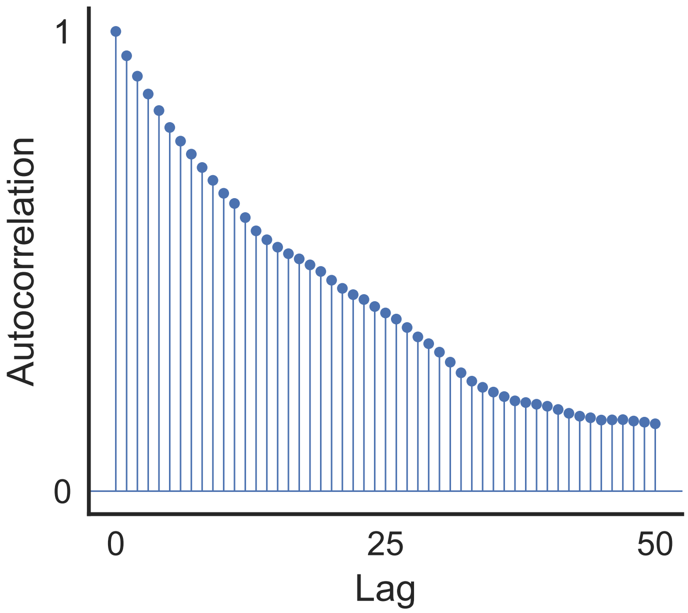


Figure 2: Autocorrelation function for TF $X_1$

Table 3: Posterior summary statistics for the inferred parameters in the single-gene motifs.

| **Parameter** | **True** | **Mean** | **SD** | $\hat{R}$ |
|---|---|---|---|---|
| $\beta_{11}$ | 10.00 | 9.075 | 0.398 | 1.00 |
| $\gamma_{11}$ | 0.00 | 0.001 | 0.467 | 1.00 |
| $\alpha_1$ | 0.05 | 0.046 | 0.002 | 1.00 |
| $\beta_{22}$ | 0.00 | −0.003 | 0.463 | 1.00 |
| $\gamma_{22}$ | 10.00 | 9.564 | 0.424 | 1.00 |
| $\alpha_2$ | 0.05 | 0.048 | 0.002 | 1.00 |

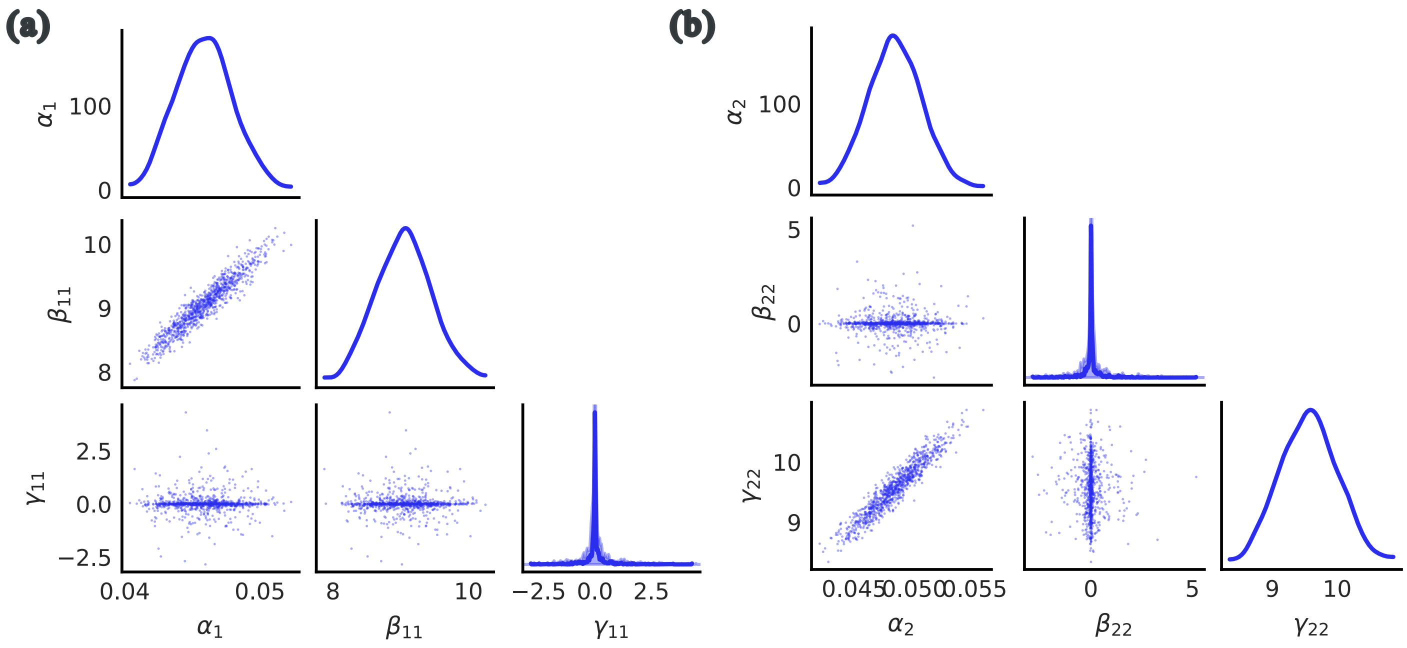


Figure 3: **Joint posterior distributions of inferred parameters for (a) self-activation and (b) self-repression motifs.**

### 3 Two-Gene Motifs

In this section, we present the data generation setup and expanded inference results for the two-gene motif with cross-regulation example introduced in the main text.

#### 3.1 Data

Synthetic time-series data was generated using the SSA. The regulatory functions and parameter values used in the simulations are summarized in Table 4.

Table 4: Propensity functions and parameter values for $X_1$ and $X_2$ in the two-gene motifs.

| Transcription Factor | Propensities | $\beta/\gamma$ | $K$ | $n$ | $\alpha$ |
|---|---|---|---|---|---|
| $X_1$ | $f_a(X_2),\ \alpha_1 X_1$ | 10 | 80 | 4 | 0.03 |
| $X_2$ | $f_r(X_1),\ \alpha_2 X_2$ | 20 | 50 | 4 | 0.02 |

#### 3.2 Definition of likelihoods

The explicit expressions for the drift and diffusion terms for each gene $X_i$ are given by:

$$\mu_1 = \left(\frac{\gamma_{11}K_1^{n_1}}{X_1^{n_1}+K_1^{n_1}} + \frac{\beta_{11}X_1^{n_1}}{X_1^{n_1}+K_1^{n_1}} + \frac{\gamma_{12}K_1^{n_1}}{X_2^{n_1}+K_1^{n_1}} + \frac{\beta_{12}X_2^{n_1}}{X_2^{n_1}+K_1^{n_1}} - \alpha_1 X_1\right)\Delta t,$$
$$\sigma_1 = \sqrt{\frac{\gamma_{11}K_1^{n_1}}{X_1^{n_1}+K_1^{n_1}} + \frac{\beta_{11}X_1^{n_1}}{X_1^{n_1}+K_1^{n_1}} + \frac{\gamma_{12}K_1^{n_1}}{X_2^{n_1}+K_1^{n_1}} + \frac{\beta_{12}X_2^{n_1}}{X_2^{n_1}+K_1^{n_1}} + \alpha_1 X_1}\sqrt{\Delta t}. \quad (2)$$

$$\mu_2 = \left( \frac{\gamma_{22} K_2^{n_2}}{X_2^{n_2} + K_2^{n_2}} + \frac{\beta_{22} X_2^{n_2}}{X_2^{n_2} + K_2^{n_2}} + \frac{\gamma_{21} K_2^{n_2}}{X_1^{n_2} + K_2^{n_2}} + \frac{\beta_{21} X_1^{n_2}}{X_1^{n_2} + K_2^{n_2}} - \alpha_2 X_2 \right) \Delta t,$$
$$\sigma_2 = \sqrt{\frac{\gamma_{22} K_2^{n_2}}{X_2^{n_2} + K_2^{n_2}} + \frac{\beta_{22} X_2^{n_2}}{X_2^{n_2} + K_2^{n_2}} + \frac{\gamma_{21} K_2^{n_2}}{X_1^{n_2} + K_2^{n_2}} + \frac{\beta_{21} X_1^{n_2}}{X_1^{n_2} + K_2^{n_2}} + \alpha_2 X_2} \sqrt{\Delta t}. \tag{3}$$

##### 3.3 Expanded Results

Fig. 4 shows box plots of all the inferred parameters, and Table 5 summarizes posterior means, standard deviations, and $\hat{R}$ values. Joint and marginal posterior distributions are shown in Fig. 5.

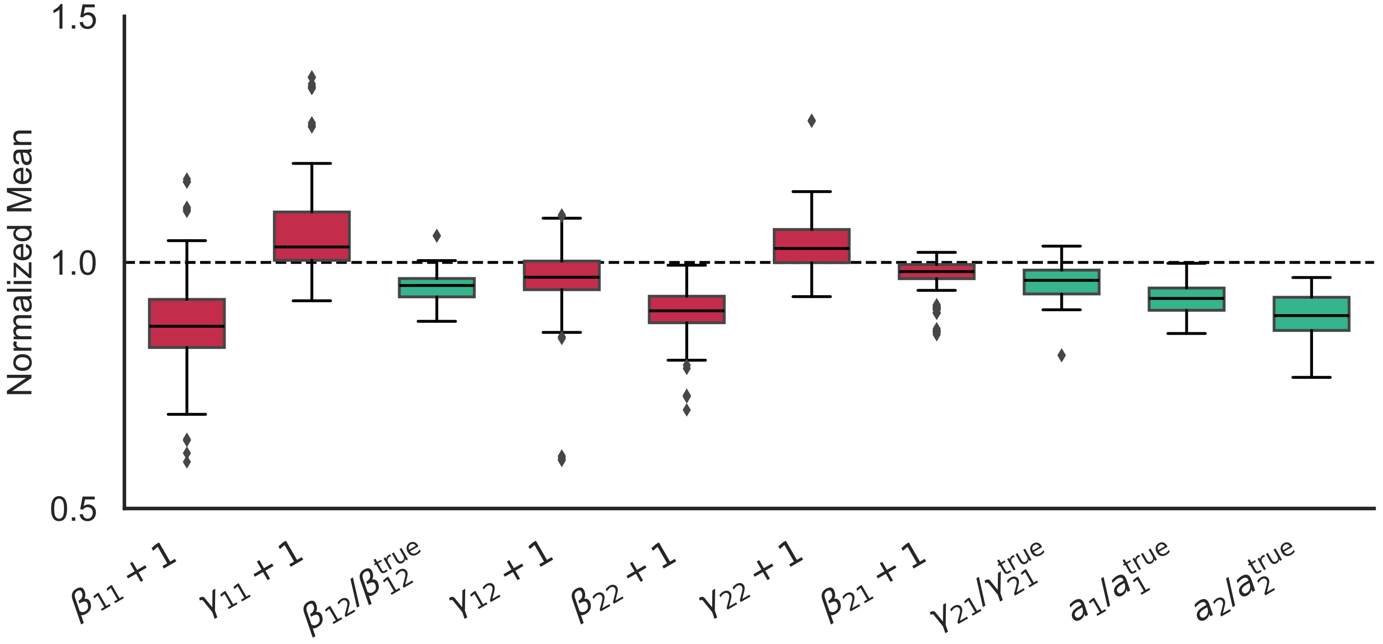


Figure 4: Box plots of all the inferred parameters for the two-gene motif.

Table 5: Posterior summaries for selected inferred parameters in the two-gene motif.

| **Parameter** | **True** | **Mean** | **SD** | $\hat{R}$ |
|---|---|---|---|---|
| $\beta_{11}$ | 0 | -0.125 | 0.474 | 1.038 |
| $\gamma_{11}$ | 0 | 0.064 | 0.261 | 1.057 |
| $\beta_{12}$ | 10 | 9.481 | 0.521 | 1.362 |
| $\gamma_{12}$ | 0 | -0.034 | 0.200 | 1.051 |
| $\beta_{21}$ | 0 | -0.026 | 0.114 | 1.046 |
| $\gamma_{21}$ | 20 | 19.183 | 1.152 | 1.255 |
| $\beta_{22}$ | 0 | -0.100 | 0.273 | 1.023 |
| $\gamma_{22}$ | 0 | 0.039 | 0.166 | 1.058 |
| $\alpha_1$ | 0.03 | 0.028 | 0.004 | 1.120 |
| $\alpha_2$ | 0.02 | 0.018 | 0.003 | 1.098 |

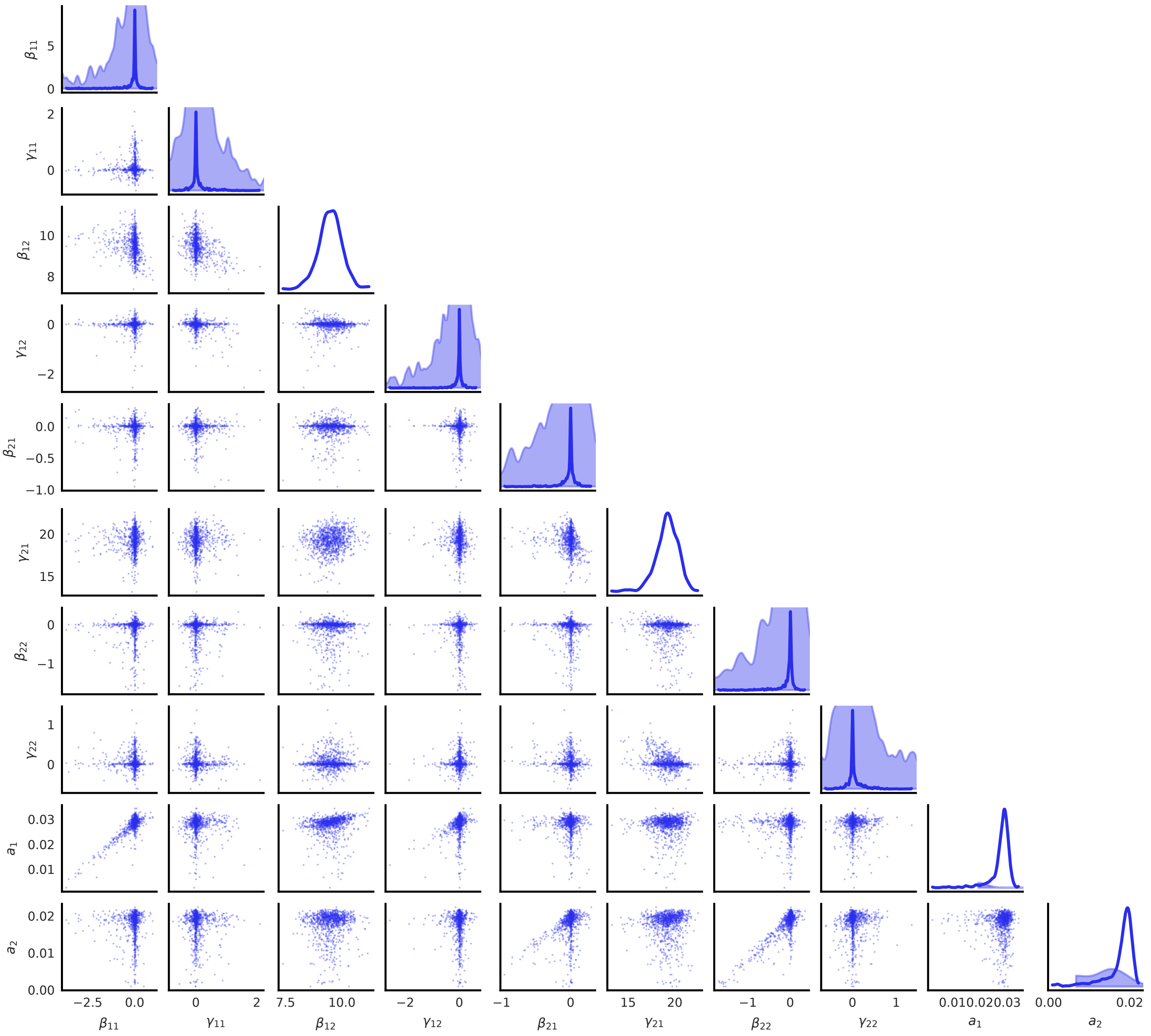

Figure 5: Joint posterior distributions of inferred parameters for the two-gene motif.

### 4 Two-Gene Motifs with an AND Gate

In this section, we provide the details for the two-gene AND-gate example described in the main text.

#### 4.1 Data

Synthetic trajectories were generated using the SSA with the propensity functions and parameter values listed in Table 6.

#### 4.2 Definition of likelihoods

The drift and diffusion terms for each TF are given by:

Table 6: Propensity functions and parameter values for $X_1$ and $X_2$ in the two-gene motif with an AND gate.

| **Transcription Factor** | **Propensities** | $\boldsymbol{\beta/\gamma/\eta}$ | $\boldsymbol{K}$ | $\boldsymbol{n}$ | $\boldsymbol{\alpha}$ |
|---|---|---|---|---|---|
| $X_1$ | $H_{rr}(X_2(t)),\ \alpha_1 X_1(t)$ | 20 | 100, 80 | 2 | 0.06 |
| $X_2$ | $f_a(X_1(t)),\ \alpha_2 X_2(t)$ | 10 | 70 | 2 | 0.055 |

$$
\begin{aligned}
\mu_1 &= \Delta t\Bigg(\eta_1\left[\frac{\omega_1\omega_2\, X_1^{n_1}X_2^{n_2} + \omega_1\tilde{\omega}_2\, X_1^{n_1}K_2^{n_2} + \tilde{\omega}_1\omega_2\, K_1^{n_1}X_2^{n_2} + \tilde{\omega}_1\tilde{\omega}_2\, K_1^{n_1}K_2^{n_2}}{(K_1^{n_1}+X_1^{n_1})(K_2^{n_2}+X_2^{n_2})}\right] \\
&\quad + \beta_{11}\frac{\theta_{11}X_1^{n_1} + \tilde{\theta}_{11}K_1^{n_1}}{X_1^{n_1}+K_1^{n_1}} + \beta_{12}\frac{\theta_{12}X_2^{n_2} + \tilde{\theta}_{12}K_2^{n_2}}{X_2^{n_2}+K_2^{n_2}} - \alpha_1 X_1\Bigg) \\
\sigma_1(t) &= \sqrt{\Delta t}\Bigg(\sqrt{\eta_1\frac{\omega_1\omega_2\, X_1^{n_1}X_2^{n_2} + \omega_1\tilde{\omega}_2\, X_1^{n_1}K_2^{n_2} + \tilde{\omega}_1\omega_2\, K_1^{n_1}X_2^{n_2} + \tilde{\omega}_1\tilde{\omega}_2\, K_1^{n_1}K_2^{n_2}}{(K_1^{n_1}+X_1^{n_1})(K_2^{n_2}+X_2^{n_2})}} \\
&\quad + \beta_{11}\frac{\theta_{11}X_1^{n_1} + \tilde{\theta}_{11}K_1^{n_1}}{X_1^{n_1}+K_1^{n_1}} + \beta_{12}\frac{\theta_{12}X_2^{n_2} + \tilde{\theta}_{12}K_2^{n_2}}{X_2^{n_2}+K_2^{n_2}} + \alpha_1 X_1\Bigg)
\end{aligned} \tag{4}
$$

$$
\begin{aligned}
\mu_2 &= \Delta t\Bigg(\eta_2\frac{\omega_3\omega_4\, X_1^{n_3}X_2^{n_2} + \omega_3\tilde{\omega}_4\, X_1^{n_3}K_2^{n_2} + \tilde{\omega}_3\omega_4\, K_3^{n_3}X_2^{n_2} + \tilde{\omega}_3\tilde{\omega}_4\, K_3^{n_3}K_2^{n_2}}{(K_3^{n_3}+X_1^{n_3})(K_2^{n_2}+X_2^{n_2})} \\
&\quad + \beta_{21}\frac{\theta_{21}X_1^{n_3} + \tilde{\theta}_{21}K_3^{n_3}}{X_1^{n_3}+K_3^{n_3}} + \beta_{22}\frac{\theta_{22}X_2^{n_3} + \tilde{\theta}_{22}K_3^{n_3}}{X_2^{n_3}+K_3^{n_3}} - \alpha_2 X_2\Bigg) \\
\sigma_2(t) &= \sqrt{\Delta t}\Bigg(\sqrt{\eta_2\frac{\omega_3\omega_4\, X_1^{n_3}X_2^{n_2} + \omega_3\tilde{\omega}_4\, X_1^{n_3}K_2^{n_2} + \tilde{\omega}_3\omega_4\, K_3^{n_3}X_2^{n_2} + \tilde{\omega}_3\tilde{\omega}_4\, K_3^{n_3}K_2^{n_2}}{(K_3^{n_3}+X_1^{n_3})(K_2^{n_2}+X_2^{n_2})}} \\
&\quad + \sqrt{\beta_{21}\frac{\theta_{21}X_1^{n_3} + \tilde{\theta}_{21}K_3^{n_3}}{X_1^{n_3}+K_3^{n_3}} + \beta_{22}\frac{\theta_{22}X_2^{n_3} + \tilde{\theta}_{22}K_3^{n_3}}{X_2^{n_3}+K_3^{n_3}} + \alpha_2 X_2}\Bigg)
\end{aligned} \tag{5}
$$

#### 4.3 Expanded Results

Table 7 summarizes the inferred parameters. All $\hat{R}$ values are approximately 1, indicating good convergence, and the posterior means are close to the true generating values. The joint posterior densities with diagonal marginals are shown in Fig. 6.

Table 7: Summary statistics for inferred parameters for 2 gene AND gate

| **Transcription Factor** | **True Value** | **Mean** | **SD** | $\boldsymbol{\hat{R}}$ |
|---|---|---|---|---|
| $\beta_{11}$ | 0 | -0.043 | 0.360 | 1.001 |
| $\beta_{12}$ | 0 | 0.088 | 0.511 | 1.000 |
| $\beta_{21}$ | 10 | 9.415 | 0.355 | 1.000 |
| $\beta_{22}$ | 0 | 0.030 | 0.386 | 1.000 |
| $\alpha_1$ | 0.06 | 0.055 | 0.002 | 1.000 |
| $\alpha_2$ | 0.055 | 0.052 | 0.002 | 1.001 |
| $\eta_1$ | 20 | 18.271 | 0.918 | 1.000 |
| $\eta_2$ | 0 | 0.024 | 0.487 | 1.000 |

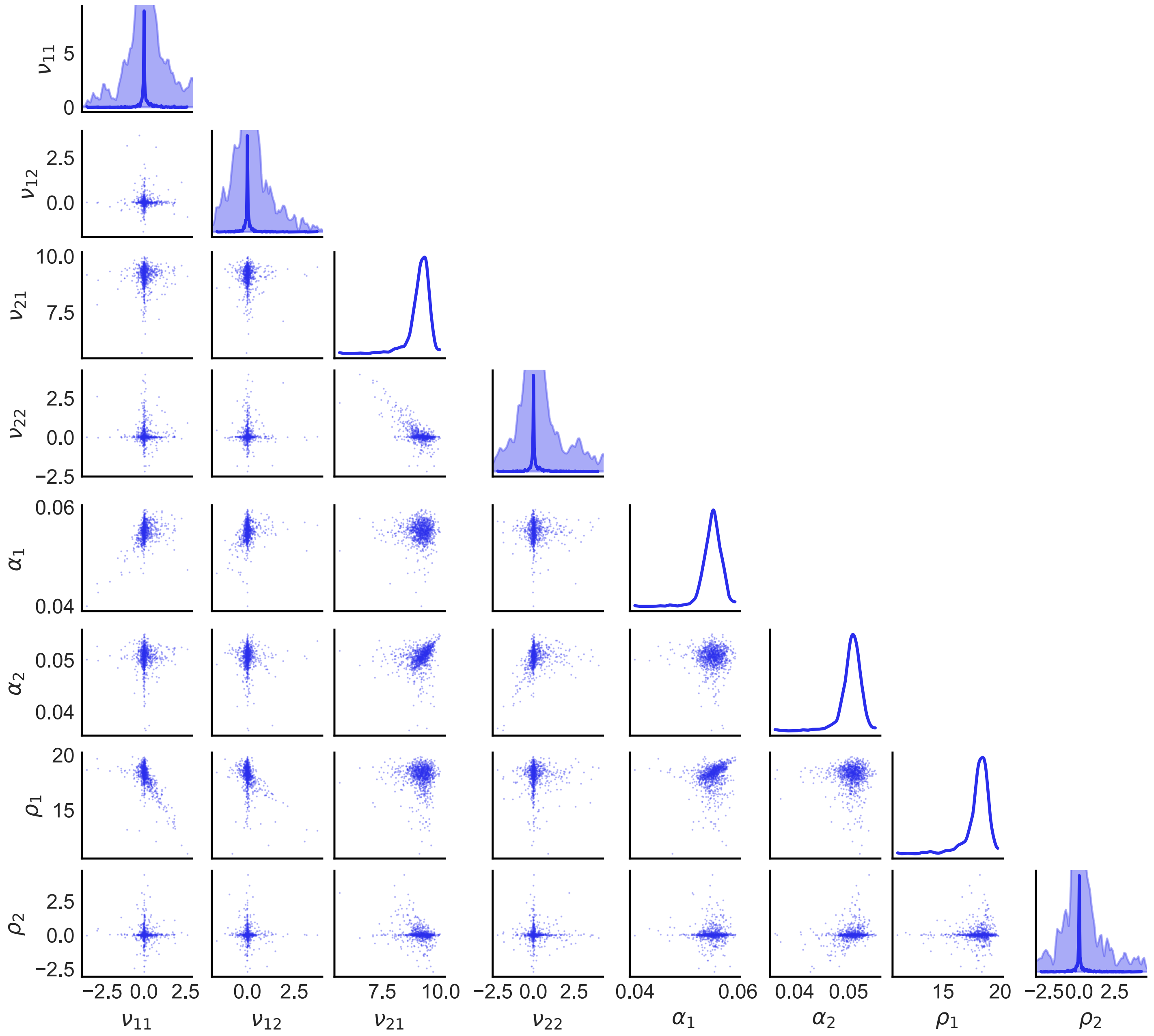


Figure 6: Joint posterior distributions for the two-gene AND-gate example.

### 5 Repressilator

In this section, we provide the full details for the three-gene repressilator example discussed in the main text.

#### 5.1 Data

Synthetic trajectories are generated using the stochastic simulation algorithm (SSA). The propensity functions and the parameter values used in the simulations for the three proteins are listed in Table 8.

Table 8: Propensity functions and parameter values for $X_1$, $X_2$, and $X_3$ in the repressilator motif.

| **Transcription Factor** | **Propensities** | $\boldsymbol{\beta/\gamma}$ | $\boldsymbol{K}$ | $\boldsymbol{n}$ | $\boldsymbol{\alpha}$ |
|---|---|---|---|---|---|
| $X_1$ | $f_r(X_3(t)),\ \alpha_1 X_1(t)$ | 10 | 90 | 2 | 0.03 |
| $X_2$ | $f_r(X_1(t)),\ \alpha_2 X_2(t)$ | 20 | 70 | 2 | 0.05 |
| $X_3$ | $f_r(X_2(t)),\ \alpha_3 X_3(t)$ | 15 | 50 | 2 | 0.04 |

#### 5.2 Definition of likelihoods

The drift and diffusion terms for each species follow directly from the general CLE-based formulation. For each $i \in \{1, 2, 3\}$,

$$\begin{aligned}
\mu_1(t) &= \Delta t\Bigg(\beta_{11}\frac{X_1^{n_1}}{K_1^{n_1}+X_1^{n_1}} + \gamma_{11}\frac{K_1^{n_1}}{K_1^{n_1}+X_1^{n_1}} + \beta_{12}\frac{X_2^{n_2}}{K_2^{n_2}+X_2^{n_2}} + \gamma_{12}\frac{K_2^{n_2}}{K_2^{n_2}+X_2^{n_2}} \\
&\qquad + \beta_{13}\frac{X_3^{n_3}}{K_3^{n_3}+X_3^{n_3}} + \gamma_{13}\frac{K_3^{n_3}}{K_3^{n_3}+X_3^{n_3}} - \alpha_1 X_1\Bigg) \\
\sigma_1(t) &= \sqrt{\Delta t}\left(\sqrt{\beta_{11}\frac{X_1^{n_1}}{K_1^{n_1}+X_1^{n_1}} + \gamma_{11}\frac{K_1^{n_1}}{K_1^{n_1}+X_1^{n_1}} + \beta_{12}\frac{X_2^{n_2}}{K_2^{n_2}+X_2^{n_2}}}\right. \\
&\qquad \left.\sqrt{+\gamma_{12}\frac{K_2^{n_2}}{K_2^{n_2}+X_2^{n_2}} + \beta_{13}\frac{X_3^{n_3}}{K_3^{n_3}+X_3^{n_3}} + \gamma_{13}\frac{K_3^{n_3}}{K_3^{n_3}+X_3^{n_3}} + \alpha_1 X_1}\right)
\end{aligned} \tag{6}$$

$$\begin{aligned}
\mu_2(t) &= \Delta t\Bigg(\beta_{22}\frac{X_2^{n_2}}{K_2^{n_2}+X_2^{n_2}} + \gamma_{22}\frac{K_2^{n_2}}{K_2^{n_2}+X_2^{n_2}} + \beta_{21}\frac{X_1(t)^{n_1}}{K_1^{n_1}+X_1^{n_1}} + \gamma_{21}\frac{K_1^{n_1}}{K_1^{n_1}+X_1^{n_1}} \\
&\qquad + \beta_{23}\frac{X_3^{n_3}}{K_3^{n_3}+X_3^{n_3}} + \gamma_{23}\frac{K_3^{n_3}}{K_3^{n_3}+X_3^{n_3}} - \alpha_2 X_2\Bigg) \\
\sigma_2(t) &= \sqrt{\Delta t}\ \sqrt{\beta_{22}\frac{X_2^{n_2}}{K_2^{n_2}+X_2^{n_2}} + \gamma_{22}\frac{K_2^{n_2}}{K_2^{n_2}+X_2^{n_2}} + \beta_{21}\frac{X_1^{n_1}}{K_1^{n_1}+X_1^{n_1}} + \gamma_{21}\frac{K_1^{n_1}}{K_1^{n_1}+X_1^{n_1}}} \\
&\qquad \sqrt{+\beta_{23}\frac{X_3^{n_3}}{K_3^{n_3}+X_3^{n_3}} + \gamma_{23}\frac{K_3^{n_3}}{K_3^{n_3}+X_3^{n_3}} + \alpha_2 X_2}
\end{aligned} \tag{7}$$

$$\begin{aligned}
\mu_3(t) &= \Delta t\Bigg(\beta_{31}\frac{X_1^{n_1}}{K_1^{n_1}+X_1^{n_1}} + \gamma_{31}\frac{K_1^{n_1}}{K_1^{n_1}+X_1^{n_1}} + \beta_{32}\frac{X_2^{n_2}}{K_2^{n_2}+X_2^{n_2}} + \gamma_{32}\frac{K_2^{n_2}}{K_2^{n_2}+X_2^{n_2}} \\
&\qquad + \beta_{33}\frac{X_3^{n_3}}{K_3^{n_3}+X_3^{n_3}} + \gamma_{33}\frac{K_3^{n_3}}{K_3^{n_3}+X_3^{n_3}} - \alpha_3 X_3\Bigg) \\
\sigma_3(t) &= \sqrt{\Delta t}\ \sqrt{\beta_{31}\frac{X_1^{n_1}}{K_1^{n_1}+X_1^{n_1}} + \gamma_{31}\frac{K_1^{n_1}}{K_1^{n_1}+X_1^{n_1}} + \beta_{32}\frac{X_2^{n_2}}{K_2^{n_2}+X_2^{n_2}} + \gamma_{32}\frac{K_2^{n_2}}{K_2^{n_2}+X_2^{n_2}}} \\
&\qquad \sqrt{+\beta_{33}\frac{X_3^{n_3}}{K_3^{n_3}+X_3^{n_3}} + \gamma_{33}\frac{K_3^{n_3}}{K_3^{n_3}+X_3^{n_3}} + \alpha_3 X_3}
\end{aligned} \tag{8}$$

#### 5.3 Expanded Results

Table 9 reports posterior summary statistics for all inferred parameters. All chains show excellent convergence ($\hat{R} \approx 1$), and the posterior means closely match the true values, indicating accurate parameter recovery. Box plots and joint posterior distributions are shown in Fig. 7 and 8, respectively.

Table 9: Posterior summary statistics with true parameter values for the Repressilator.

| **Parameter** | **True Value** | **Mean** | **SD** | $\hat{R}$ |
|---|---|---|---|---|
| $\beta_{11}$ | 0 | 0.003 | 0.218 | 1.00 |
| $\beta_{12}$ | 0 | 0.114 | 0.279 | 1.00 |
| $\beta_{13}$ | 0 | 0.016 | 0.160 | 1.00 |
| $\beta_{21}$ | 0 | 0.013 | 0.127 | 1.00 |
| $\beta_{22}$ | 0 | -0.007 | 0.211 | 1.00 |
| $\beta_{23}$ | 0 | 0.087 | 0.230 | 1.00 |
| $\beta_{31}$ | 0 | 0.010 | 0.147 | 1.00 |
| $\beta_{32}$ | 0 | -0.017 | 0.135 | 1.00 |
| $\beta_{33}$ | 0 | -0.042 | 0.287 | 1.00 |
| $\gamma_{11}$ | 0 | 0.125 | 0.417 | 1.00 |
| $\gamma_{12}$ | 0 | -0.052 | 0.290 | 1.00 |
| $\gamma_{13}$ | 10 | 9.485 | 0.536 | 1.00 |
| $\gamma_{21}$ | 20 | 18.287 | 0.909 | 1.00 |
| $\gamma_{22}$ | 0 | 0.048 | 0.203 | 1.00 |
| $\gamma_{23}$ | 0 | -0.032 | 0.195 | 1.00 |
| $\gamma_{31}$ | 0 | -0.049 | 0.353 | 1.00 |
| $\gamma_{32}$ | 15 | 14.197 | 0.524 | 1.00 |
| $\gamma_{33}$ | 0 | 0.011 | 0.158 | 1.00 |
| $\alpha_1$ | 0.03 | 0.030 | 0.001 | 1.00 |
| $\alpha_2$ | 0.05 | 0.046 | 0.002 | 1.00 |
| $\alpha_3$ | 0.04 | 0.038 | 0.002 | 1.00 |

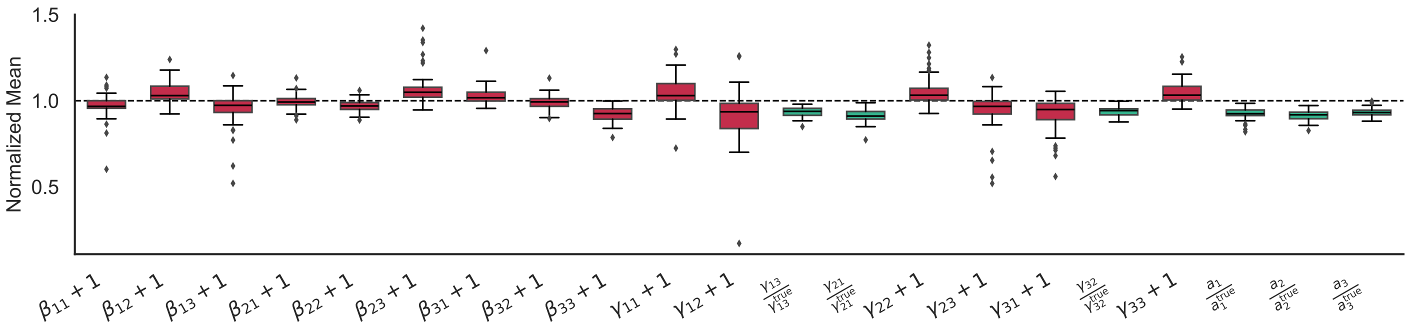


Figure 7: Box plots of posterior parameter means for the Repressilator.

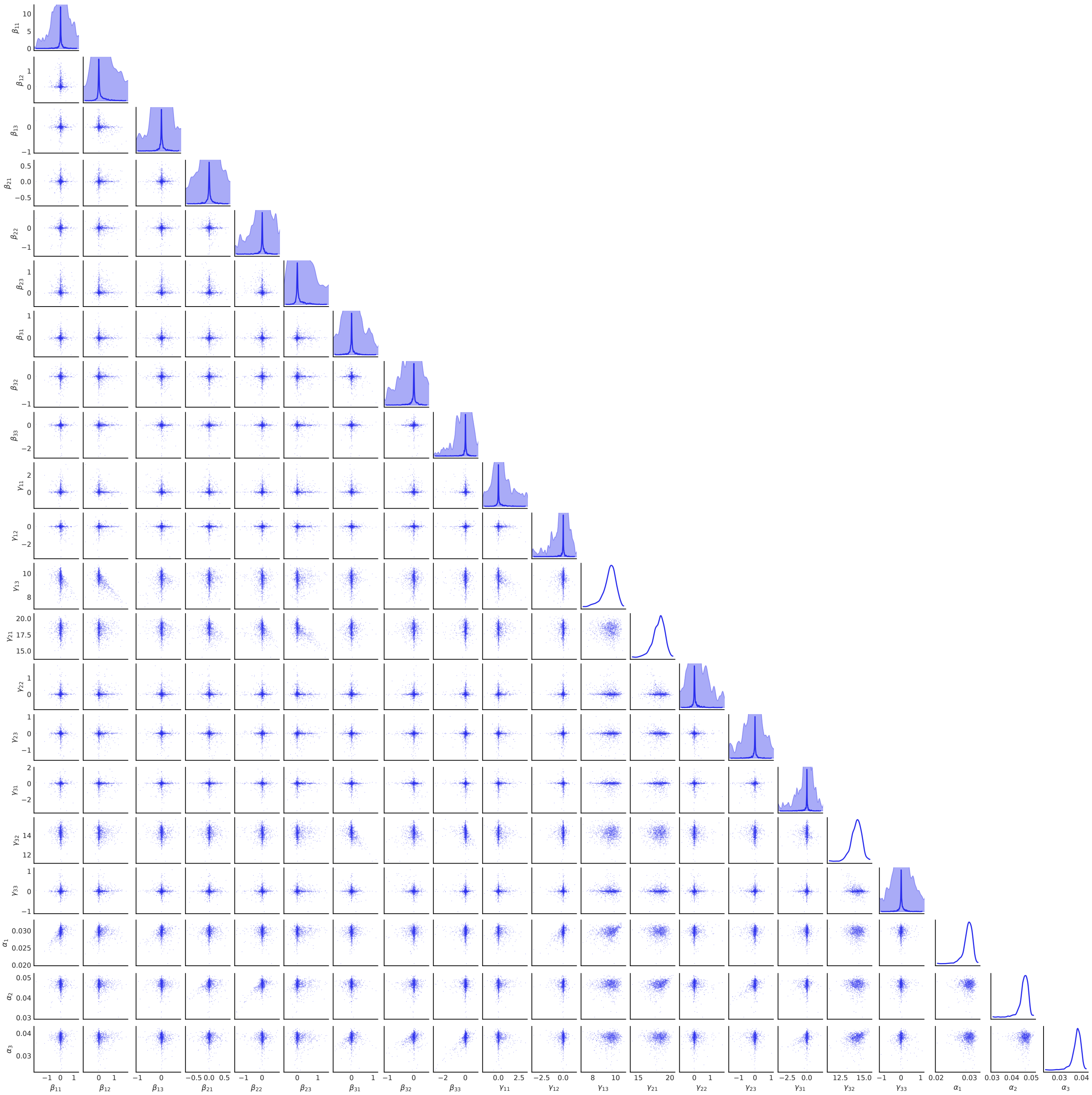

Figure 8: Joint posterior distributions for the Repressilator parameters.

### 6 Coherent Type 1 Motif

We now provide details for the Coherent Type 1 feed-forward loop discussed in the main text. This motif consists of a single regulatory interaction from $X_1$ to $X_2$, and an AND-gate at $X_3$ driven jointly by $X_1$ and $X_2$. Inference is performed in two steps: in Step 1, we infer the network topology using the model in Eq. (9); in Step 2, we use the inferred topology as prior structural knowledge and estimate all kinetic parameters using the full model in Eq. (8).

#### 6.1 Data

Table 10 reports the propensity functions and parameter values used to generate the SSA data.

Table 10: Propensity functions and parameter values for $X_1$, $X_2$, and $X_3$ in the Coherent Type 1 motif.

| **Transcription Factor** | **Propensities** | $\boldsymbol{\beta/\gamma/\eta}$ | $\boldsymbol{K}$ | $\boldsymbol{n}$ | $\boldsymbol{\alpha}$ |
|---|---|---|---|---|---|
| $X_1$ | $k = 5,\ \alpha_1 X_1$ | – | – | 3 | 0.06 |
| $X_2$ | $f_a(X_1),\ \alpha_2 X_2$ | 20 | 150 | 2 | 0.045 |
| $X_3$ | $H_{aa}(X_1, X_2),\ \alpha_3 X_3$ | 20 | 10, 50 | 3 | 0.05 |

#### 6.2 Definition of likelihoods

Inference proceeds in two steps. First, we provide the drift and diffusion terms used in Step 1 for topology inference, followed by the corresponding expressions used in Step 2 for parameter estimation.

##### 6.2.1 Step 1

$$
\begin{aligned}
\mu_1 = \Delta t \Bigg( & \nu_{12}\left[\theta_{12}\frac{X_2^n}{X_2^n+K^n} + \hat{\theta}_{12}\frac{K^n}{X_2^n+K^n}\right] + \nu_{13}\left[\theta_{13}\frac{X_3^{n_1}}{X_3^{n_1}+K_1^{n_1}} + \hat{\theta}_{13}\frac{K_1^{n_1}}{X_3^{n_1}+K_1^{n_1}}\right] \\
& + \rho_1\Bigg[\omega_{12}\omega_{13}\frac{X_2^n X_3^{n_1}}{(X_2^n+K^n)(X_3^{n_1}+K_1^{n_1})} + \hat{\omega}_{12}\omega_{13}\frac{K^n X_3^{n_1}}{(X_2^n+K^n)(X_3^{n_1}+K_1^{n_1})} \\
& \quad + \omega_{12}\hat{\omega}_{13}\frac{X_2^n K_1^{n_1}}{(X_2^n+K^n)(X_3^{n_1}+K_1^{n_1})} + \hat{\omega}_{12}\hat{\omega}_{13}\frac{K^n K_1^{n_1}}{(X_2^n+K^n)(X_3^{n_1}+K_1^{n_1})}\Bigg] + r_1 - \alpha_1 X_1\Bigg) \\
\sigma_1 = \sqrt{\Delta t}\, & \sqrt{\nu_{12}\left[\theta_{12}\frac{X_2^n}{X_2^n+K^n} + \hat{\theta}_{12}\frac{K^n}{X_2^n+K^n}\right] + \nu_{13}\left[\theta_{13}\frac{X_3^{n_1}}{X_3^{n_1}+K_1^{n_1}} + \hat{\theta}_{13}\frac{K_1^{n_1}}{X_3^{n_1}+K_1^{n_1}}\right]} \\
& + \sqrt{\rho_1\left[\omega_{12}\omega_{13}\frac{X_2^n X_3^{n_1}}{(X_2^n+K^n)(X_3^{n_1}+K_1^{n_1})} + \hat{\omega}_{12}\omega_{13}\frac{K^n X_3^{n_1}}{(X_2^n+K^n)(X_3^{n_1}+K_1^{n_1})}\right.} \\
& \quad + \sqrt{\left.\omega_{12}\hat{\omega}_{13}\frac{X_2^n K_1^{n_1}}{(X_2^n+K^n)(X_3^{n_1}+K_1^{n_1})} + \hat{\omega}_{12}\hat{\omega}_{13}\frac{K^n K_1^{n_1}}{(X_2^n+K^n)(X_3^{n_1}+K_1^{n_1})}\right] + r_1 + \alpha_1 X_1}
\end{aligned}
\tag{9}
$$

$$\mu_2(t) = \Delta t \Bigg( \nu_{21} \left[ \theta_{21} \frac{X_1^n}{X_1^n + K^n} + \hat{\theta}_{21} \frac{K^n}{X_1^n + K^n} \right] + \nu_{23} \left[ \theta_{23} \frac{X_3^{n_2}}{X_3^{n_2} + K_2^{n_2}} + \hat{\theta}_{23} \frac{K_2^{n_2}}{X_3^{n_2} + K_2^{n_2}} \right]$$

$$+ \rho_2 \Bigg[ \omega_{21}\omega_{23} \frac{X_1^n X_3^{n_2}}{(X_1^n + K^n)(X_3^{n_2} + K_2^{n_2})} + \hat{\omega}_{21}\omega_{23} \frac{K^n X_3^{n_2}}{(X_1^n + K^n)(X_3^{n_2} + K_2^{n_2})}$$
$$+ \omega_{21}\hat{\omega}_{23} \frac{X_1^n K_2^{n_2}}{(X_1^n + K^n)(X_3^{n_2} + K_2^{n_2})} + \hat{\omega}_{21}\hat{\omega}_{23} \frac{K^n K_2^{n_2}}{(X_1^n + K^n)(X_3^{n_2} + K_2^{n_2})} \Bigg] + r_2 - \alpha_2 X_2 \Bigg)$$

$$\sigma_2(t) = \sqrt{\Delta t} \sqrt{\nu_{21} \left[ \theta_{21} \frac{X_1^n}{X_1^n + K^n} + \hat{\theta}_{21} \frac{K^n}{X_1^n + K^n} \right] + \nu_{23} \left[ \theta_{23} \frac{X_3^{n_2}}{X_3^{n_2} + K_2^{n_2}} + \hat{\theta}_{23} \frac{K_2^{n_2}}{X_3^{n_2} + K_2^{n_2}} \right]} \tag{10}$$

$$\sqrt{+\rho_2 \Bigg[ \omega_{21}\omega_{23} \frac{X_1^n X_3^{n_2}}{(X_1^n + K^n)(X_3^{n_2} + K_2^{n_2})} + \hat{\omega}_{21}\omega_{23} \frac{K^n X_3^{n_2}}{(X_1^n + K^n)(X_3^{n_2} + K_2^{n_2})}}$$
$$\sqrt{+\omega_{21}\hat{\omega}_{23} \frac{X_1^n K_2^{n_2}}{(X_1^n + K^n)(X_3^{n_2} + K_2^{n_2})} + \hat{\omega}_{21}\hat{\omega}_{23} \frac{K^n K_2^{n_2}}{(X_1^n + K^n)(X_3^{n_2} + K_2^{n_2})} \Bigg] + \alpha_2 X_2 + r_2}$$

$$\mu_3 = \Delta t \Bigg( \nu_{31} \frac{\theta_{31} X_1^{n_1} + \hat{\theta}_{31} K_1^{n_1}}{X_1^{n_1} + K_1^{n_1}} + \nu_{32} \frac{\theta_{32} X_2^{n_2} + \hat{\theta}_{32} K_2^{n_2}}{X_2^{n_2} + K_2^{n_2}}$$

$$+ \rho_3 \Bigg[ \omega_{31}\omega_{32} \frac{X_1^{n_1} X_2^{n_2}}{(X_1^{n_1} + K_1^{n_1})(X_2^{n_2} + K_2^{n_2})} + \hat{\omega}_{31}\omega_{32} \frac{K_1^{n_1} X_2^{n_2}}{(X_1^{n_1} + K_1^{n_1})(X_2^{n_2} + K_2^{n_2})}$$
$$+ \omega_{31}\hat{\omega}_{32} \frac{X_1^{n_1} K_2^{n_2}}{(X_1^{n_1} + K_1^{n_1})(X_2^{n_2} + K_2^{n_2})} + \hat{\omega}_{31}\hat{\omega}_{32} \frac{K_1^{n_1} K_2^{n_2}}{(X_1^{n_1} + K_1^{n_1})(X_2^{n_2} + K_2^{n_2})} \Bigg] + r_3 - \alpha_3 X_3 \Bigg)$$

$$\sigma_3 = \sqrt{\Delta t} \sqrt{\nu_{31} \frac{\theta_{31} X_1^{n_1} + \hat{\theta}_{31} K_1^{n_1}}{X_1^{n_1} + K_1^{n_1}} + \nu_{32} \frac{\theta_{32} X_2^{n_2} + \hat{\theta}_{32} K_2^{n_2}}{X_2^{n_2} + K_2^{n_2}}} \tag{11}$$

$$\sqrt{+\rho_3 \Bigg[ \omega_{31}\omega_{32} \frac{X_1^{n_1} X_2^{n_2}}{(X_1^{n_1} + K_1^{n_1})(X_2^{n_2} + K_2^{n_2})} + \hat{\omega}_{31}\omega_{32} \frac{K_1^{n_1} X_2^{n_2}}{(X_1^{n_1} + K_1^{n_1})(X_2^{n_2} + K_2^{n_2})}}$$
$$\sqrt{+\omega_{31}\hat{\omega}_{32} \frac{X_1^{n_1} K_2^{n_2}}{(X_1^{n_1} + K_1^{n_1})(X_2^{n_2} + K_2^{n_2})} + \hat{\omega}_{31}\hat{\omega}_{32} \frac{K_1^{n_1} K_2^{n_2}}{(X_1^{n_1} + K_1^{n_1})(X_2^{n_2} + K_2^{n_2})} \Bigg] + \alpha_3 X_3 + r_3}$$

**Step2**

$$\mu_1 = \Delta t \left( r_1 - \alpha_1 X_1 \right)$$

$$\sigma_1 = \sqrt{\Delta t} \sqrt{r_1 + \alpha_1 X_1} \tag{12}$$

$$
\mu_2 = \Delta t\left(\beta_{21}\frac{X_1^n}{X_1^n + K^n} + \gamma_{21}\frac{K^n}{X_1^n + K^n} + \beta_{23}\frac{X_3^{n_2}}{X_3^{n_2} + K_2^{n_2}} + \gamma_{23}\frac{K_2^{n_2}}{X_3^{n_2} + K_2^{n_2}} + r_2 - \alpha_2 X_2\right)
$$

$$
\sigma_2 = \sqrt{\Delta t}\sqrt{\beta_{21}\frac{X_1^n}{X_1^n + K^n} + \gamma_{21}\frac{K^n}{X_1^n + K^n} + \beta_{23}\frac{X_3^{n_2}}{X_3^{n_2} + K_2^{n_2}} + \gamma_{23}\frac{K_2^{n_2}}{X_3^{n_2} + K_2^{n_2}} + \alpha_2 X_2 + r_2} \tag{13}
$$

$$
\begin{aligned}
\mu_3 = \Delta t\Bigg(&\eta_1\frac{X_1^{n_1}X_2^{n_2}}{(X_1^{n_1} + K_1^{n_1})(X_2^{n_2} + K_2^{n_2})} + \eta_2\frac{K_1^{n_1}X_2^{n_2}}{(X_1^{n_1} + K_1^{n_1})(X_2^{n_2} + K_2^{n_2})} \\
&+ \eta_3\frac{X_1^{n_1}K_2^{n_2}}{(X_1^{n_1} + K_1^{n_1})(X_2^{n_2} + K_2^{n_2})} + \eta_4\frac{K_1^{n_1}K_2^{n_2}}{(X_1^{n_1} + K_1^{n_1})(X_2^{n_2} + K_2^{n_2})} + r_3 - \alpha_3 X_3\Bigg)
\end{aligned} \tag{14}
$$

$$
\begin{aligned}
\sigma_3 = \sqrt{\Delta t}&\sqrt{\eta_1\frac{X_1^{n_1}X_2^{n_2}}{(X_1^{n_1} + K_1^{n_1})(X_2^{n_2} + K_2^{n_2})} + \eta_2\frac{K_1^{n_1}X_2^{n_2}}{(X_1^{n_1} + K_1^{n_1})(X_2^{n_2} + K_2^{n_2})}} \\
&\sqrt{+\eta_3\frac{X_1^{n_1}K_2^{n_2}}{(X_1^{n_1} + K_1^{n_1})(X_2^{n_2} + K_2^{n_2})} + \eta_4\frac{K_1^{n_1}K_2^{n_2}}{(X_1^{n_1} + K_1^{n_1})(X_2^{n_2} + K_2^{n_2})} + \alpha_3 X_3 + r_3}
\end{aligned}
$$

#### 6.3 Expanded Results

**Step 1.** Fig. 9 shows the joint posterior distributions obtained in the 2nd inference step. Table 11 reports the posterior summaries for all parameters in Step 1. Presence of the activation/Repression or both is inferred through $\theta$ and $\omega$ values. And the parameters inferred close to 0 are pruned.

**Step 2.** Using the inferred topology from Step 1, we perform the second inference step with the reduced parameter space. Table 12 summarizes the posterior distributions, showing tight credible intervals and accurate recovery of the true parameters.

#### 6.4 Identifiability Considerations

To illustrate the identifiability issues that motivate the two-step inference procedure, Fig. 10 shows regions of parameter space where different combinations of interaction strengths yield nearly indistinguishable trajectories.

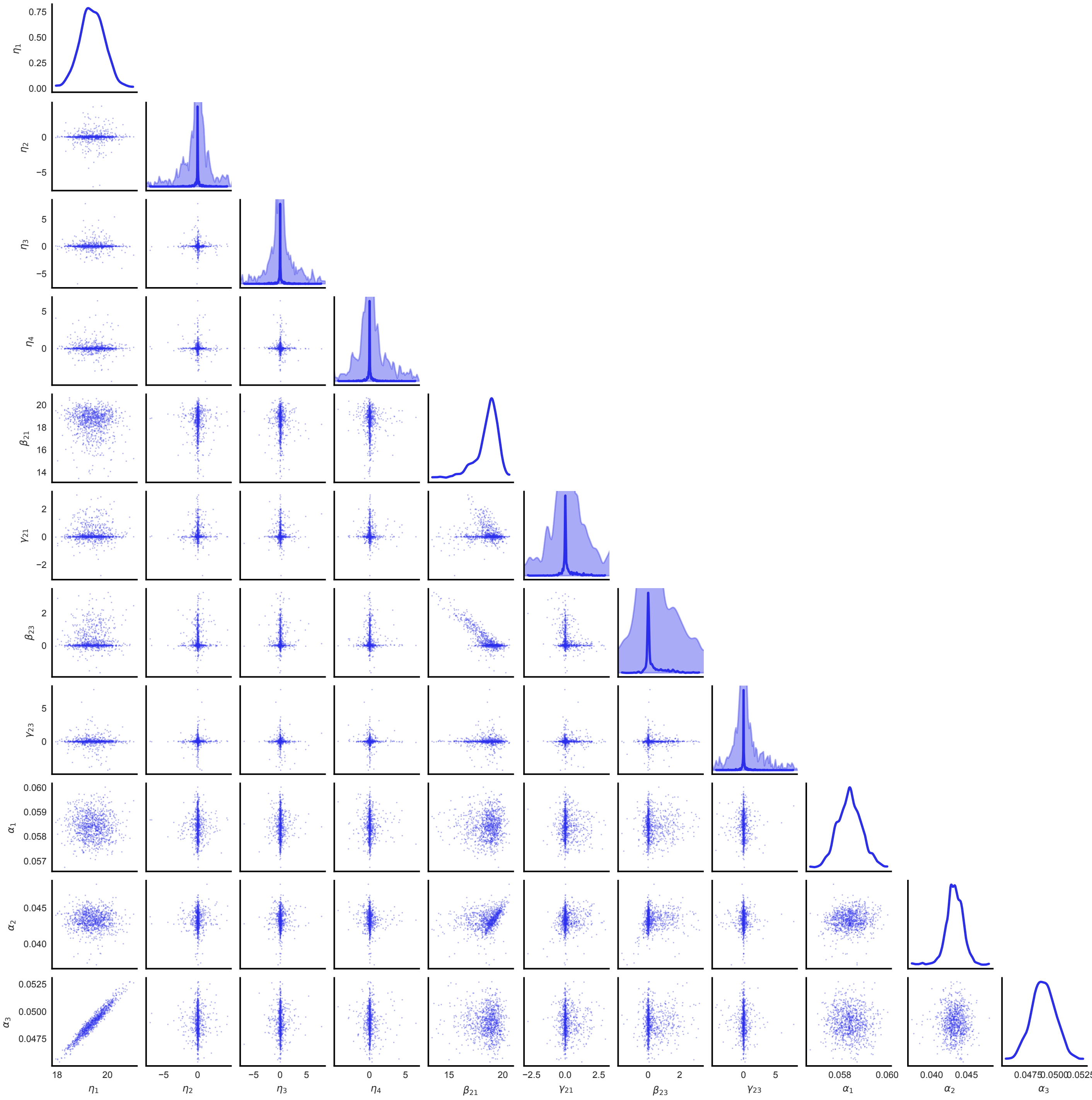


Figure 9: Joint posterior distributions obtained in the 2nd inference step

Table 11: Posterior summary statistics for Step 1 of the Coherent Type 1 motif.

| **Parameter** | **True** | **Mean** | **SD** | $\hat{R}$ |
|---|---|---|---|---|
| $\rho_1$ | 0 | -0.055 | 0.391 | 1.000 |
| $\rho_2$ | 0 | 7.752 | 8.979 | 1.013 |
| $\rho_3$ | 20 | 6.798 | 8.895 | 1.024 |
| $\nu_{12}$ | 0 | -0.062 | 0.275 | 1.002 |
| $\nu_{13}$ | 0 | -0.035 | 0.359 | 1.001 |
| $\nu_{21}$ | 20 | 10.731 | 8.971 | 1.013 |
| $\nu_{23}$ | 0 | 0.027 | 0.398 | 1.005 |
| $\nu_{31}$ | 0 | 4.366 | 7.740 | 1.004 |
| $\nu_{32}$ | 0 | 7.841 | 9.131 | 1.010 |
| $\theta_{12}$ | 1 | 0.844 | 0.362 | 1.001 |
| $\theta_{13}$ | 1 | 0.828 | 0.377 | 1.001 |
| $\theta_{21}$ | 1 | 0.952 | 0.214 | 1.002 |
| $\theta_{23}$ | 1 | 0.839 | 0.368 | 1.001 |
| $\theta_{31}$ | 1 | 0.883 | 0.321 | 1.001 |
| $\theta_{32}$ | 1 | 0.916 | 0.277 | 1.002 |
| $\omega_{12}$ | 1 | 0.845 | 0.362 | 1.001 |
| $\omega_{13}$ | 1 | 0.834 | 0.372 | 1.000 |
| $\omega_{21}$ | 1 | 0.907 | 0.291 | 1.002 |
| $\omega_{23}$ | 1 | 0.903 | 0.296 | 1.002 |
| $\omega_{31}$ | 1 | 0.893 | 0.309 | 1.001 |
| $\omega_{32}$ | 1 | 0.892 | 0.310 | 1.001 |

Table 12: Posterior summary statistics for Step 2 inference.

| **Parameter** | **True** | **Mean** | **SD** | $\hat{R}$ |
|---|---|---|---|---|
| $\eta_1$ | 20 | 19.395 | 0.497 | 1.00 |
| $\eta_2$ | 0 | 0.003 | 0.725 | 1.00 |
| $\eta_3$ | 0 | -0.007 | 0.730 | 1.00 |
| $\eta_4$ | 0 | 0.001 | 0.725 | 1.00 |
| $\beta_{21}$ | 20 | 18.528 | 1.124 | 1.00 |
| $\gamma_{21}$ | 0 | 0.157 | 0.507 | 1.00 |
| $\beta_{23}$ | 0 | 0.296 | 0.637 | 1.00 |
| $\gamma_{23}$ | 0 | 0.005 | 0.740 | 1.00 |
| $\alpha_1$ | 0.06 | 0.058 | 0.000 | 1.00 |
| $\alpha_2$ | 0.045 | 0.043 | 0.001 | 1.00 |
| $\alpha_3$ | 0.05 | 0.049 | 0.001 | 1.00 |

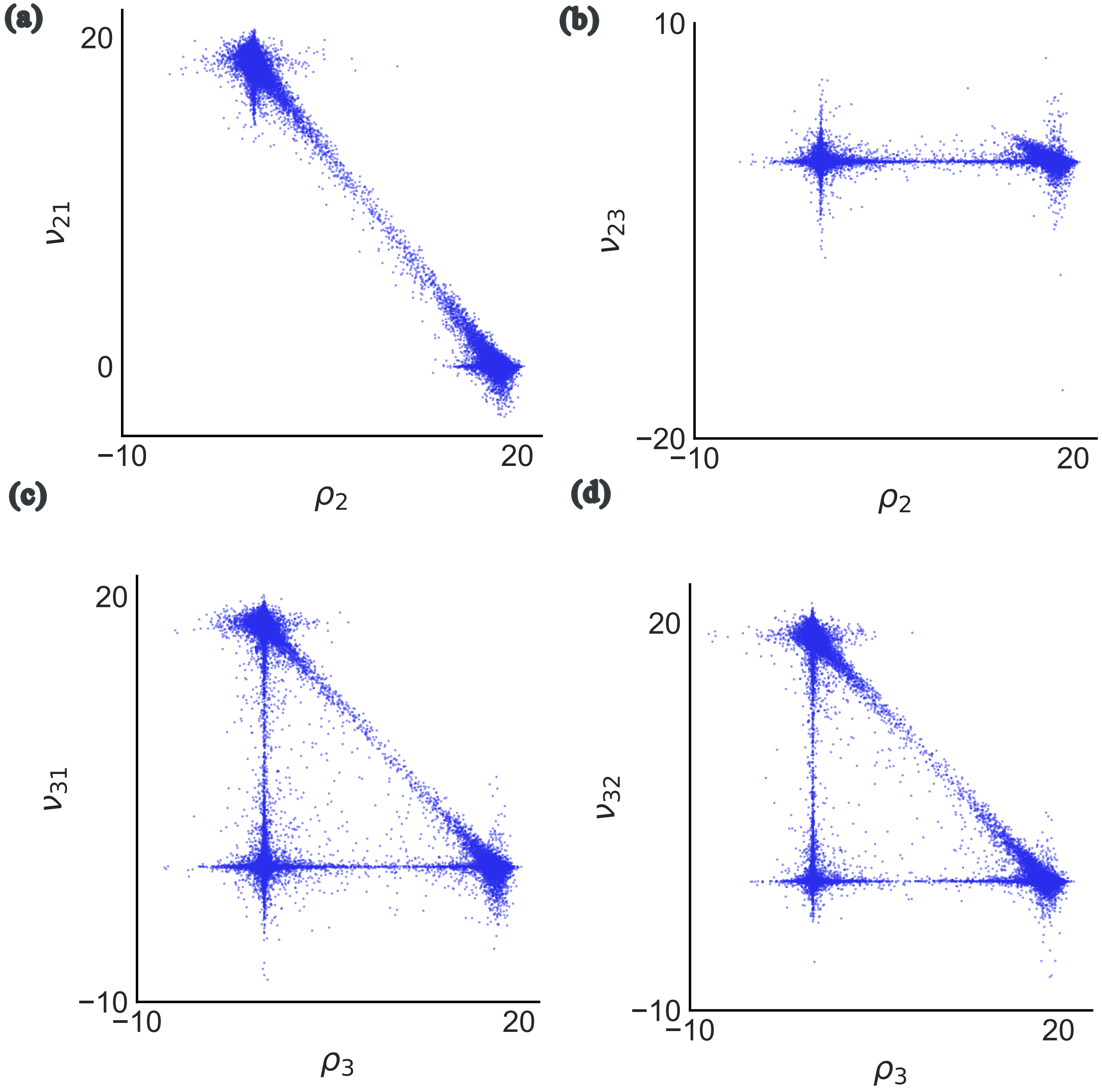


Figure 10: Illustration of identifiability issues for the Coherent Type 1 motif for step 1 of the inference procedure.
